# Cortical Origins of the Flash-Lag Effect distortions: The Influence of Retinotopic Map Architecture

**DOI:** 10.1101/2025.09.15.676318

**Authors:** Salvatore Giancani, Sandrine Chemla, Frédéric Chavane, Manuel Vidal

## Abstract

The flash-lag effect (FLE) is an illusion whereby the position of a moving object is perceived as being offset in the direction of movement when compared to a flashed static stimulus. This perceptual misalignment has been posited as a key phenomenon in explaining our ability to accurately predict the future position of moving objects, despite the delays in neuronal processing. Our working hypothesis is that the FLE is resulting from the anticipation generated by propagation of neural activity within visual cortical retinotopical maps. According to this hypothesis, the FLE should be affected by discontinuities and anisotropies of the retinotopic map architecture. Using psychophysics we show that the FLE is strongly affected by crossing and the direction of motion in respect to retinotopic features, such as vertical and horizontal meridians and the fovea. The specificity of how early visual cortical retinotopic maps are splitted and magnified around these features led us to suggest that the FLE distortions emerge from propagation in retinotopically organized networks, particularly V1. This work bridges the gap between human psychophysics and the known constraints of retinotopic maps layout, offering a testable framework for future studies of motion position encoding in the visual hierarchy.

## Introduction

The perception of the position of a flashing object often lags behind the perceived position of a moving stimulus. This is a phenomenon first described by Mackay (1958) and later termed the “flash-lag effect” (FLE) by Nijhawan (1994) that linked this effect to a compensatory phenomenon of the delay for the transmission of visual information. In other words, the FLE would be a consequence of the brain’s anticipatory processing of motion, named “motion extrapolation” (Hogendoorn 2020; Turner et al. 2023). Since Nijhawan (1994), the effect has been extensively studied using various stimuli, including single and double moving bars, rotating annuli, rotating objects, and dots in apparent motion (Hubbard 2014; Eagleman and Sejnowski 2007; Nagai, Kazai, and Yagi 2002; Li, Shim, and Cavanagh 2014). The effect has been explored in different positions of the field of view (varying both in azimuth and elevation), motion directions (horizontal, vertical), speeds, contrasts, colors, and timing of the flashing stimulus appearing whether at the start (flash-initiated condition or Fröhlich effect (Fröhlich 1923)), mid-point, or end of the moving stimulus trajectory (Hubbard 2014; Eagleman and Sejnowski 2007). It has been claimed that the misalignment between moving and flashed objects increases with the uncertainty of the moving object position (Kanai et al. 2004): the FLE increases for example, at lower contrasts or at high speeds of the moving stimulus (for these and other examples of position estimations confound see Hubbard 2014). Since the re-discovery by Nijhawan (1994), the scientific community has debated on the phenomenological explanation of the effect: the main theoretical debates regards the brain’s predictive versus postdictive nature and its relationship to phenomenal awareness (Eagleman and Sejnowski 2000; 2007; Nijhawan 2002; Hubbard 2014; Hogendoorn 2020). However, our understanding of the physiological mechanics of the phenomenon stays limited, and only recently, efforts have been made to address this knowledge gap (Turner, Sexton, and Hogendoorn 2023).

Numerous studies (Hogendoorn and Burkitt 2018; Johnson et al. 2023; Turner et al. 2024; 2025) demonstrated consistent motion extrapolation in humans’ EEG. However, the aforementioned works do not point out precisely to a neural structure responsible for the motion extrapolation, indicating generically the visual pathway as actor, suggesting a cumulative effect that adds up at each neural processing step. Some fMRI studies on humans demonstrated that the encoding of the neural position of a moving stimulus is shifted (Schneider et al. 2019; Kohler et al. 2017; Maus et al. 2013; Whitney et al. 2003). Nevertheless, some of the studies showed differences in position encoding for early and not higher visual areas (Whitney et al. 2003; Kohler et al. 2017) and some others only for higher level areas (Maus, Fischer, and Whitney 2013; Schneider et al. 2019). The neural origin of the FLE thus remains highly debated. However, the evidence gathered over decades on non-human models suggests that motion extrapolation may result from the cumulative contributions of multiple visual processing levels. In Berry et al. (1999) it was shown on salamander and rabbit’s retinas that a moving bar generates an anticipatory response within the receptive field of retinal ganglion cells (RGC) through a gain control mechanism. A different mechanism has been suggested in the primary visual cortex of anesthetized cats (Jancke et al. 2004), in anesthetized (Guo et al. 2007) and awake non-human primates (NHP) (Subramaniyan et al. 2018; Benvenuti et al. 2020), implying cortico-cortical propagations and anticipatory activity arising before reaching the receptive field of cortical neurons.

One possible mechanism for modulating the encoding of static versus moving object positions is connectivity between neurons whose receptive fields (RFs) do not overlap. There are two main types: inter-cortical feedback from higher areas (Figure1A cyan) and intra-cortical horizontal connectivity (Figure1A blue). These connections, which can be seen as deviations from the retinotopic organization, carry information about the presence of a stimulus in the receptive field periphery and hereby mediate lateral interactions. Importantly, these are mostly confined to subthreshold modulation (Bringuier et al. 1999) or modulation of concomitant activation to the center (Angelucci and Bressloff 2006). For a moving stimulus, these cortico-cortical connections might provide converging information about the stimulus’ arrival in neurons’ RFs before those neurons receive direct feedforward input from the stimulus entering the RF. This can only occur if the latency to propagate information along these cortico-cortical routes (L_H_ or L_FB_, Figure1A) is shorter than the overall delay to receive information from the direct feedforward pathway (L _H_ or L_FB_ < L_bar_). The latter is determined by the time for the object to reach the RF position plus the retino-thalamo-cortical latency. In primates, both feedback and horizontal connections can meet this condition for stimulus speeds below 100-200°/s. Fast-propagating myelinated inter-cortical axons (Girard et al. 2001) with a high degree of divergence in both feedforward and feedback directions (Salin et al. 1989; Angelucci et al. 2002; Kennedy and Bullier 1985) can convey information about several degrees around the receptive field in less than 10-20 ms, albeit with little retinotopic precision. In contrast, horizontal intra-cortical axons are mostly non-myelinated, with slower conduction velocities (≈0.3–0.5 m/s) (Grinvald et al. 1994; Bringuier et al. 1999; Girard et al. 2001; Reynaud et al. 2012; Muller et al. 2014), but due to retino-cortical magnification, their conduction still corresponds to equivalent visual motion in the same ∼100– 200°/s range. Intra-cortical propagation has the advantage of keeping precise retinotopic information since they occur between neurons with small RF along a retinotopic map with high resolution, like V1 (Lee et al. 1998).

Intra-cortical propagation within cortical retinotopic maps could therefore generate a reduction in neuronal latency, potentially contributing to the Flash Lag Effect (FLE), we refer to this as the *Propagation within Retinotopic Map* (PRM) hypothesis. This hypothesis is supported by converging evidences from cat or NHP, in anesthetized or awake state, indicating that both the direction and the length of a stimulus’ trajectory modulate neural activity within the retinotopic map (Jancke et al. 2004; Guo et al. 2007; Subramaniyan et al. 2018; Benvenuti et al. 2020) . Notably, using single-units, voltage-sensitive dye imaging and LFP recordings, Benvenuti et al. (2020) revealed that the latency of the evoked response to a moving bar gradually decreases as the bar follows a longer trajectory before reaching the RF. Interestingly, the PRM hypothesis yields strong, two testable predictions: first, any discontinuity of the retinotopic map should strongly affect the FLE, and second, any increase/decrease of the magnification factor along the retinotopic maps should diminish/enhance the FLE, respectively. For the first case, a key feature of cortical retinotopic architecture is the splitting of the visual field by the visual meridians. The vertical meridian (VM), separating the left and right visual fields, is represented at the V1-V2 boundary, with each hemisphere processing the contralateral field (see Figure 1B, white arrow). Neurons encoding the VM are linked via callosal projections through the corpus callosum, enabling interhemispheric communication (Hubel and Wiesel 1967; Innocenti et al. 2022). The horizontal meridian (HM), dividing the upper and lower visual fields, does not form an anatomical discontinuity in V1 but does so in downstream areas, such as V2 and V3 (Shipp et al. 1995; Yu et al. 2019; Ribeiro et al. 2023, Figure 1B gray arrow), with the upper and lower visual fields represented ventrally and dorsally. Hence, an anatomical discontinuity in the retinotopic map of all visual areas characterize the VM representation, while an anatomical discontinuity for areas downstream from V1 characterize the HM. The first prediction was tested by Benvenuti et al (2020) by measuring the effect of crossing the VM on the reduction of latency of V1 population response. The latency reduction observed for bars moving within contralateral hemifields disappears when the bars originate from the ipsilateral hemifield and cross the VM, where connectivity involves interhemispheric callosal pathways instead of intracortical horizontal connections. This suggests that the reduction in latency results from the propagation of activity within, rather than between, retinotopic maps. Thus, intracortical connectivity is the most likely candidate for the anticipatory activation observed in the neural data of Benvenuti et al (2020). The second prediction relates to the non-linear projection of the visual field on the cortical space, e.g. in V1 the visual field follows a log-polar transformation (Schwartz 1977; Schira, Wade, and Tyler 2007). This transformation produces a disproportionate representation of the fovea, which covers only 1% of the retinal surface but takes 50% of the full visual cortex (Vater et al. 2022), and approximately 30% of V1’s cortical surface (Yu, Chaplin, and Rosa 2015). We therefore expect that the FLE for a stimulus moving at constant velocity should decrease when going towards the fovea.

**Figure 1.**
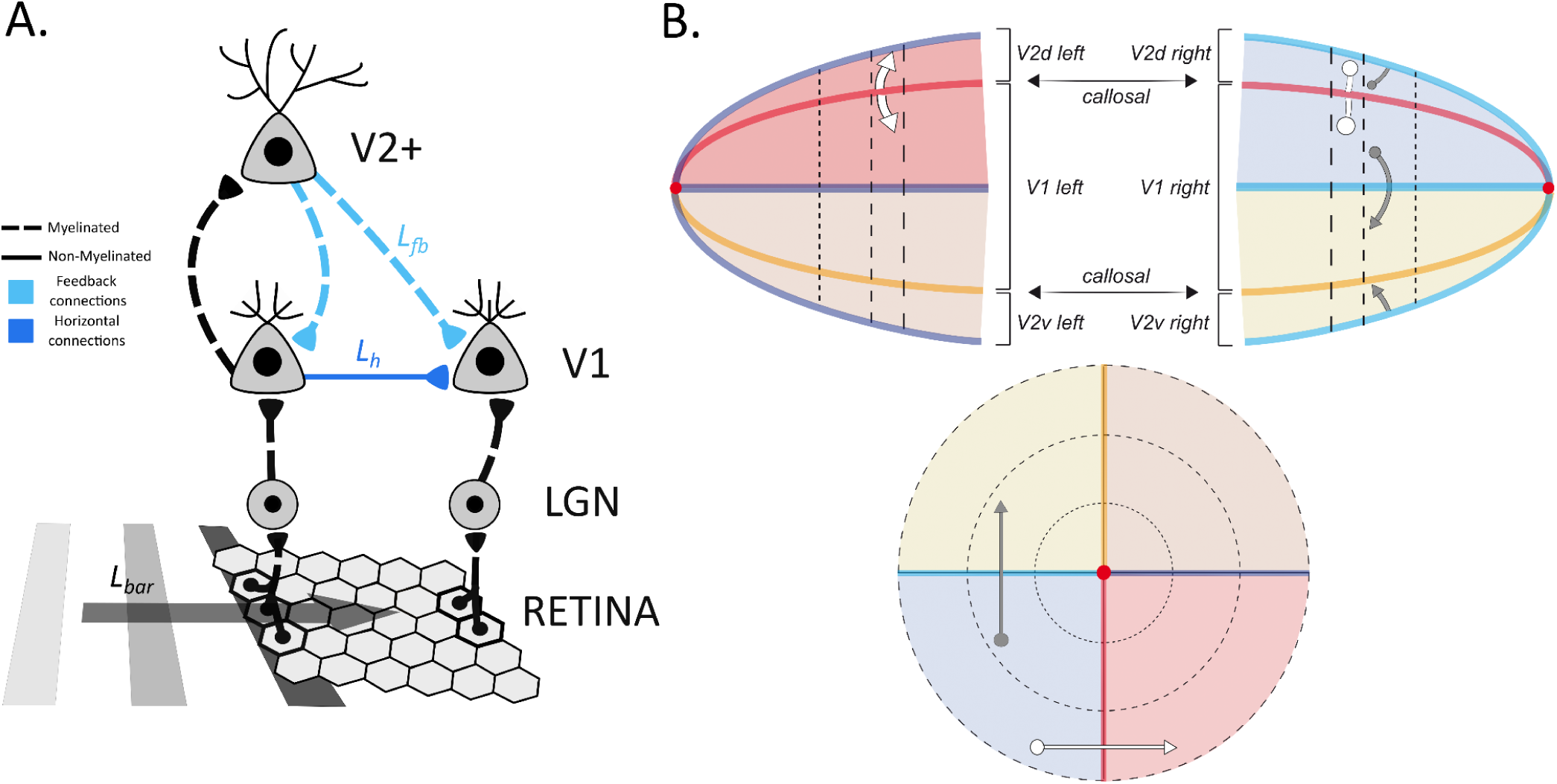
Illustration of the Propagation within Retinotopic Map (PRM) hypothesis. **(A)** Schematic of the propagation of information within a retinotopic map. A moving bar activates the retina and takes some time to reach the trajectory endpoint (*L_bar_*, gray). In a sequence of feedforward activations, mediated by myelinated fibers (black dashed lines), to the lateral geniculate nucleus (LGN) and the primary visual area (V1). At this level, two different but concurrent types of propagation can occur: the information passes to higher visual areas (V2+) through feedforward myelinated fibers and returns to V1 with fast and myelinated feedback connections (light blue dashed lines) with high spatial divergence and reaching the representation of the trajectory endpoint in V1 with latency *L_fb_*(cyan); in parallel, the information propagates within V1 through slower horizontal connectivity ( blue line) and activates V1 representation of the trajectory endpoint with a latency *L_h_*. Both dynamics may induce a drop in temporal latency if *L_h_* or *L_lb_* < *L_bar_*, mediating an early activation of cortical areas otherwise still untouched by the feedforward flow of the bar’s movement. **(B)** Schematic of the visual field with color-coded retinotopic features and quadrants. In the lower panel, a representation of the visual field is shown, with evenly spaced dashed isoeccentric circular lines and bold colored isopolar lines, corresponding to horizontal (HM, cyan and purple for left and right HM respectively) and vertical meridians (VM, orange and red for upper and lower VM respectively). The crossing of HM and VM represents the fovea (red dot). The lower hemifields are represented in light red and light blue (respectively right and left), and the upper hemifields are represented in light orange and light yellow (respectively right and left). A horizontal white arrow crosses the lower VM, while a vertical grey arrow crosses the left HM. In the upper panel, flat maps of V1 and V2 are displayed. The dashed isoeccentric circular lines are now represented as vertical lines and no longer equispaced due to the rescaling caused by the cortical magnification factor. At the outermost part of the two schematized hemispheres is the foveal representation (the red dots). Right and left hemispheres are separated along the V1/V2 border in the dorsal and ventral positions for lower (red) and upper (orange) VM respectively. Neurons close to the border are communicating through callosal connections (black arrows) that bind the right and left hemifields through the vertical meridian. It follows that the cortical representation of the white arrow crossing the lower VM appears split along the cortical representation of the lower VM. The representation of the arrow’s head and tail is located on the dorsal left and right hemifield, both for V1 and V2. The cortical representation of V2+ is splitted into quadrants, with left and right hemifield represented contralaterally and upper and lower hemifield represented in dorsal and ventral V2. As a result, the V2 cortical representation of the vertical grey arrow that crosses the horizontal meridian in the right hemifield has its head and tail represented in the ventral and dorsal parts of V2, respectively, generating a continuous activation in V1.

Here we used psychophysics to investigate the FLE and designed stimuli at specific spatial configurations to test the two predictions arising from the PRM hypothesis. Our results show a high FLE when both the bar horizontal motion trajectory and the flashed reference remain within the same hemifield, an effect that drops suddenly just after the moving bar crosses the vertical meridian (VM) where the flash was presented. This effect does not occur when the stimulus moves vertically toward the fovea, crossing the horizontal meridian (HM). Instead, we observe a slight increase of the FLE after crossing the HM and a higher FLE for downward motion. Finally, we designed the experiment to independently manipulate eccentricity and motion direction relative to the fovea, horizontal meridian (HM), and vertical meridian (VM), allowing us to tease apart their respective contributions to the FLE. By applying a geometric modeling approach, we quantified the contribution of each factor to the observed effect. Our results revealed that eccentricity and motion direction toward or away from the VM had the strongest impact on FLE magnitude, exceeding the influence of motion direction relative to the fovea or HM.

## Results

Our working hypothesis is that the flash-lag effect arises from the intra-cortical propagation of information within visual retinotopic maps, a framework we term the PRM hypothesis. According to this view, the FLE should be modulated by both discontinuities in the retinotopic layout and changes in cortical magnification, including its anisotropic distribution across vertical and horizontal dimensions (Himmelberg, Winawer, et al. 2023). These structural factors are expected to influence our ability to represent the position of moving stimuli ahead of their actual locations (Figure 1).

To test this hypothesis, we conducted a series of experiments in which we measured the strength of the FLE at different spatial locations of the visual field and for various motion directions (e.g. horizontal, vertical or oblique). Participants had to compare the perceived position of a bar translating at constant speed relative to a briefly flashed single/pair of squares aligned with the bar’s orientation (see Amadeo et al. 2022; Wang, Reynaud, and Hess 2021; Kanai, Sheth, and Shimojo 2004).

### Experiment 1

In a first experiment, we tested how the FLE changes when crossing the VM and whether its strength depends on the traveled distance. Participants were presented with bars translating horizontally below the HM (right-to-left, speed of 6.6°/s and distance ranging from 1° to 4°) before a square was flashed above the bar with a given horizontal offset. The task was to judge whether the bar was perceived to the right or to the left of the square (Figure 2A). The flash offset was varied using a Bayesian adaptive method to estimate the participants point of subjective equivalence and the standard deviation (PSE and SD), i.e. the position of the square where it is perceived as horizontally aligned with the bar and its precision. We compared the FLE observed in two different test positions (the spatial location of the bar at the time the square is flashed), either 0.5° before the VM (*same* hemifield, Figure 2B blue) or 0.5° after the VM (*crossing* hemifield, red). The first result was a dramatic global reduction of 0.28° of the FLE measured just after crossing the VM compared to just before (0.45° *vs.* 0.17°, t(9)=-8.04, p<0.0001), and this reduction was consistent across the 4 tested traveled distances (all p<0.001). The distance traveled had no influence: motion past history did not increase nor decrease the FLE observed. We conducted two control experiments to test if these findings depend on the motion direction (i.e. different FLE for bars moving leftward than rightward), or on the tested hemifield (i.e. different FLE for bars moving rightward in the left hemifield and leftward in the right hemifield). We found a similar reduction of the FLE after crossing the VM (0.39° *vs.* 0.14°, t(7)=-3.92, p<0.005) when reversing the motion direction, and no difference between hemifields (0.44° *vs.* 0.45°, t(9)=0.23, p=0.8) within the same range of distances (see supplementary Figure 1). As a consequence, in the next experiment we kept only rightward. Results for single subjects are shown in supplementary figures 2, 3, and 4.

**Figure 2.**
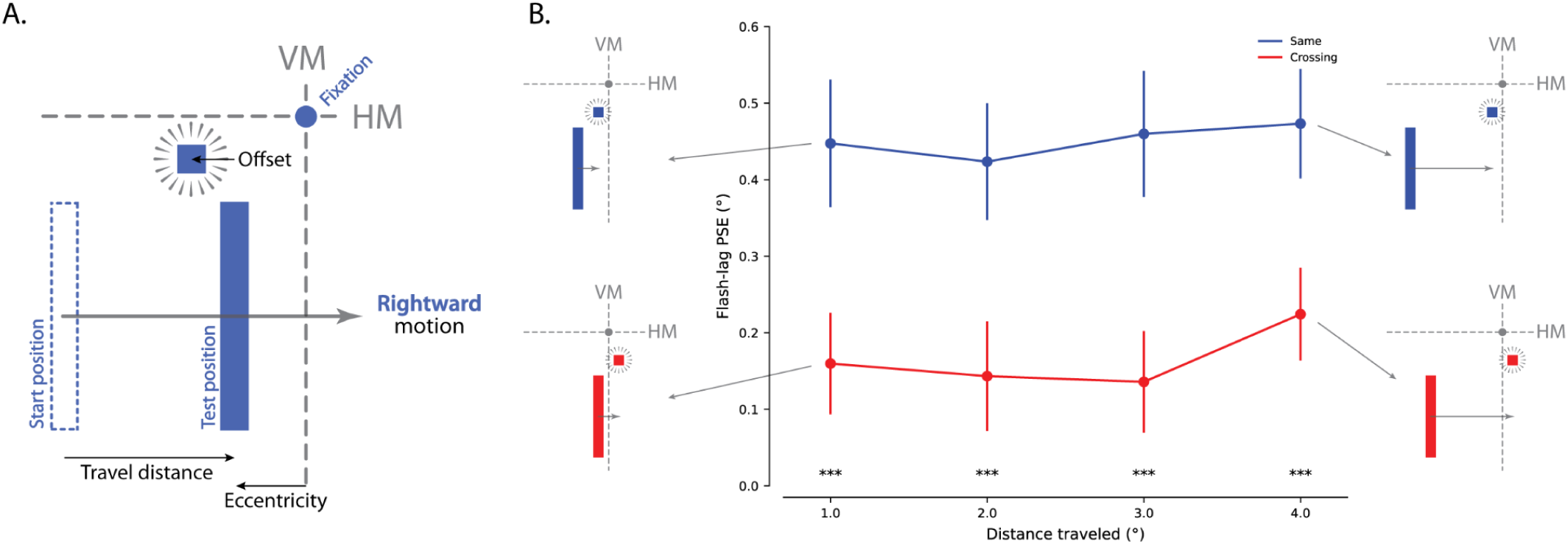
Experiment 1: The FLE is selective to the hemfield where the motion occurs. **(A)** Schematic of the flash-lag visual task and glossary of the manipulated experimental factors. The bar is moving rightward, with the start position in the lower-left hemifield. The distance between the starting position and the test position corresponds to the traveled distance, with both starting and test positions that can have different eccentricities. The square flashes with a certain offset in respect to the bar test position. The dashed lines represent the vertical (VM) and horizontal meridians (HM). The small blue disk represents the fovea (and fixation point). **(B)** The strength of the flash-lag effect (FLE) in degrees measured 0.5° before the VM (*same* hemifield, blue dots) or 0.5° after the VM (*crossing* hemifield, red dots) is represented as a function of the distance traveled (1° to 4°). Dots and error bars represent the mean ± standard error of PSE across subjects. Illustrations of the stimuli used for both conditions in the 1° and 4° distance are shown on the side.

### Experiment 2

The second experiment was designed to characterize precisely the drop of the FLE observed after crossing the VM. In this experiment, the bar moved along the HM for a distance of 2° when a pair of squares -above and below the bar-was flashed with a given horizontal offset. The horizontal eccentricity of the test position ranged from -2° to +2° in steps of 0.5° (Figure 3), so that, the bar motion was either going toward (VM-petal, -2° to -0.5°, blue) or away from the VM (VM-fugal, +0.5° to +2°, red and green) at the moment of the flash. Consistent with the findings of the first experiment, the mean FLE measured when the bar remained in the same hemifield (-2° to -0.5°, blue) was significantly larger than when crossing the VM (+0.5° to +2°, red), with 0.38° and 0.05°, respectively (t(9)=6.59, p<0.0001). Within the VM zone (-0.5° to +0.5°), the FLE dropped linearly with a slope of -0.34 (°s of PSE per °s of visual eccentricity, p<0.03). These results reveal that the drastic reduction occurs within a sharp range of tested eccentricities. Finally, there was only a limited rebuild of the FLE after crossing the VM (green) that was not significant, as it only occurred in 4 participants (see supplementary Figure 5, 6 and 7). Interestingly, the estimated SD of the PSE increases for larger positive eccentricities, indicating that the task becomes increasingly harder to perform.

**Figure 3.**
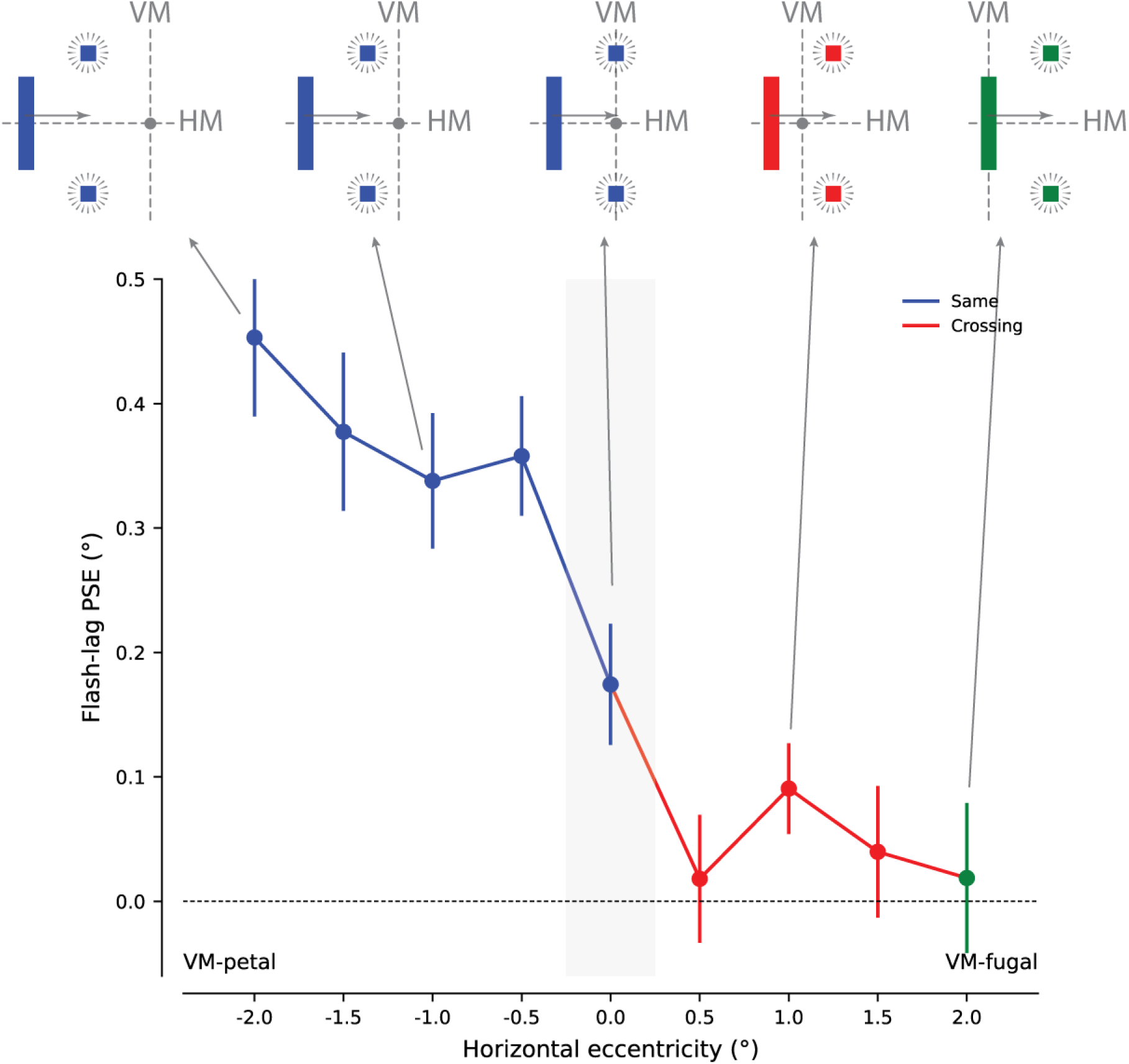
Experiment 2: The FLE drops very rapidly when crossing the VM. The strength of the flash-lag effect (FLE) in degrees as a function of the horizontal eccentricity of the tested position. We distinguish three scenarios (represented by the three colors): when the bar moves toward the VM (VM-petal direction) and the flashing squares occurs before crossing the VM (in blue); when the bar crosses the VM (VM-petal then -fugal directions) before the flash occurs (in red) and finally when the bar starts at the fovea and moves entirely in VM-fugal direction (in green). Spatially coherent illustrations of the stimuli for selected conditions are visible on the top of the figure. The gray shaded area around 0° eccentricity highlights the range where the 0.5° bar width overlaps the VM.

### Experiment 3

To further test the role of anatomical discontinuities of retinotopic maps, the third experiment was designed to characterize how the FLE evolves after crossing the HM. In this experiment, a horizontal bar was translated vertically along the VM, either moving upward or downward, for a distance of 2° when a pair of squares - right and left of the bar - was flashed at a given vertical offset. The vertical eccentricity of the test position defining the conditions are given relative to the motion direction, so that we can compare the upward and downward directions seamlessly. Four vertical eccentricities relative to motion were compared: -2°, -0.5°, +0.5° and +2° (Figure 4). At the moment of the flash, the bar was moving either toward the HM for negative eccentricities (HM-petal, blue) or away from the HM for positive eccentricities (HM-fugal, red and green). Contrarily to what was reported when crossing the VM (experiment 1 and 2), at 0.5° eccentricity, the FLE was higher when the bar crossed the HM (HM-fugal in red) than when it remained in the same hemifield (HM-petal in blue), for both upward (0.19° vs 0.06°, t(9)=3.29, p<0.01) and downward motions (0.34° vs 0.16°, t(9)=3.26, p<0.01). At 2° eccentricity, the FLE was not different between HM-petal and HM-fugal whether moving upward or downward (p>0.5 for both). Finally, upward and downward motion directions generated significantly different FLE at +0.5° (t(9)=3.59, p<0.006) and nearly at -0.5° (t(9)=2.02, p=0.075). Hence, contrary to the VM crossing and at small eccentricities, the FLE was selective to the motion direction, being stronger for bars moving downward compared to upwards. Single subjects results showed in supplementary figures 8, 9 and 10.

**Figure 4.**
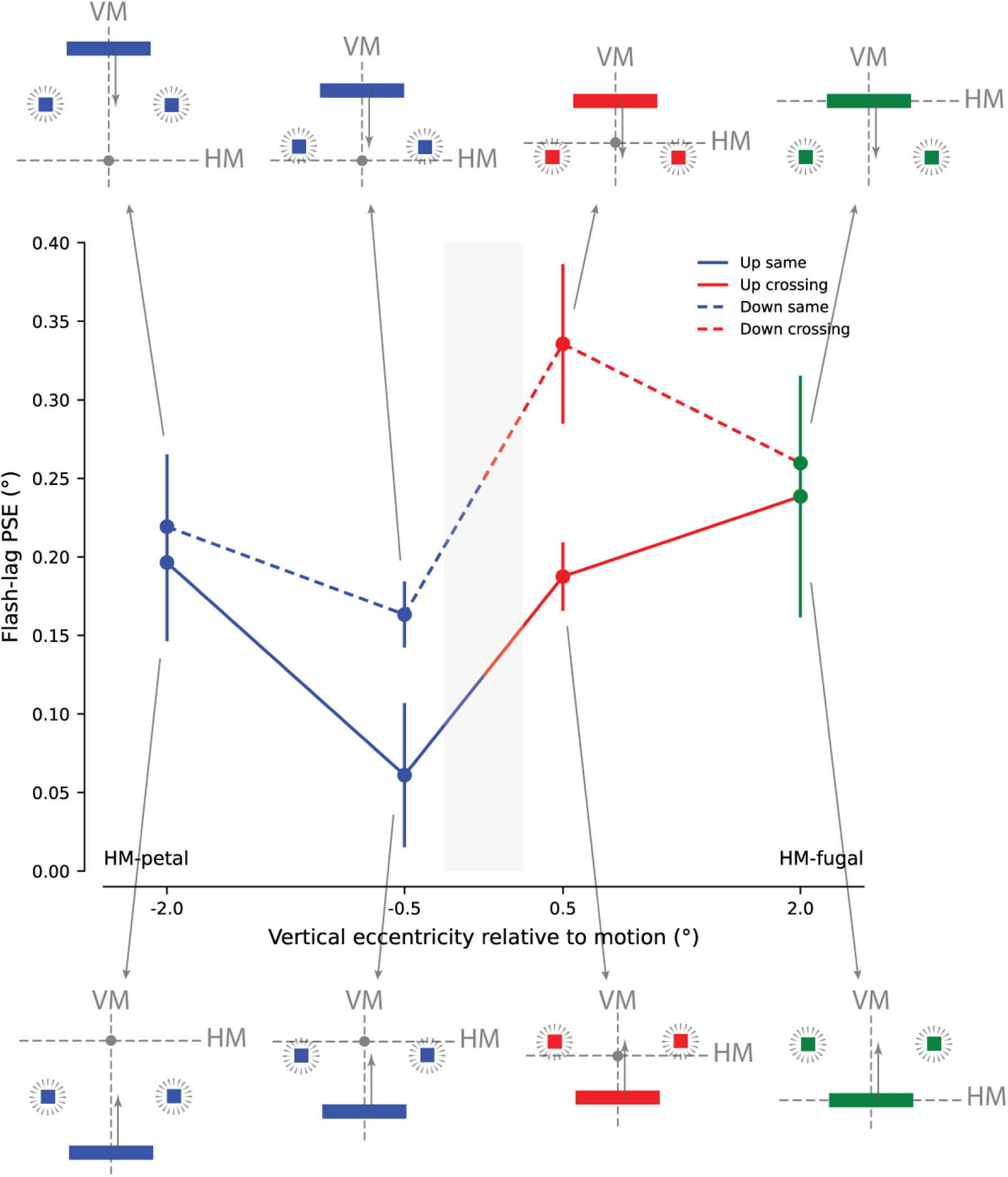
Experiment 3: The FLE around the HM. The strength of the flash-lag effect (FLE) in degrees as a function of the vertical eccentricity relative to motion of the tested position. The bar could move upward (solid line) or downward (dashed line). We distinguish three scenarios (represented by the three colors): when the bar moves toward the HM and the flashing test occurs before crossing - bar moving in HM-petal (in blue) -; the flashing test occurs after the bar crosses the HM(in red) - and finally when the bar starts to move from fovea and then flashing test occurs - bar moving entirely in HM-fugal direction (in green) -. Illustrations of the stimuli are visible on the top of the figure. The gray shaded area around 0° eccentricity highlights the range where the 0.5° bar width overlaps the HM.

From the three previous experiments, we came to the conclusion that other retinotopic features could affect the FLE. First, the FLE reduction was stronger in Experiment 2 compared to Experiment 1. Given that the main difference between these experiments was that only the VM was crossed in Experiment 1, whereas both VM and fovea were crossed in Experiment 2, it suggests that the fovea could also play a role. Furthermore, in Experiment 2, we noticed that the FLE was much stronger for motion towards the VM (blue) than away from the VM (red and green). In Experiment 3, crossing the HM modulates FLE differently than crossing the VM, pointing to a distinct functional role for the HM.

### Experiment 4

The fourth experiment was designed to fully disentangle the physiological biases on the FLE introduced by visual motion toward and away from the fovea, the VM or the HM. In this experiment, we measured the FLE in 4 test positions of the visual field for 8 motion directions (0°, 45°, 90°, 135°, 180°, 225°, 270°, 315°) resulting in a total of 32 conditions. The test position was either foveal ([0°, 0°]), on the VM but away from the fovea ([0°, -2°]), on the HM but away from the fovea ([2°, 0°]), or away from the HM, VM and fovea ([+2°, -2°]). As in previous experiments, the bar moved 2° before two squares appeared above and below the bar, at a variable offset (see Bayesian Adaptive Method in Methods). In this experiment, the bar did not cross VM, HM or fovea (see Supplementary 11A for a visual schematic). A first clear effect is the tested position in the visual field, with the conditions at position [2°, -2°] showing a larger FLE modulation than other positions (Figure 5A). A second effect is the motion direction, except for the foveal position, [0°, 0°] where the FLE remains around 0.2° for all eight directions. In all the other positions ([+2°, -2°], [0°, -2°] and [+2°, 0°]) we observe an increase of FLE in foveo-petal vs foveo-fugal motions, with values near 0.35° for foveo-petal and slightly below 0.2° for foveo-fugal. This difference is only significant at position [2°, -2°] (0.48° vs 0.08°, t(9)=3.73, p=0.006). Consistently, the diagonal motion directions at positions [2°, -2°] and [+2°, 0°] also show a stronger FLE when they contain a motion-component towards the VM (directions 135° and 225°) compared to away from the VM (direction 45° and 315°).

**Figure 5.**
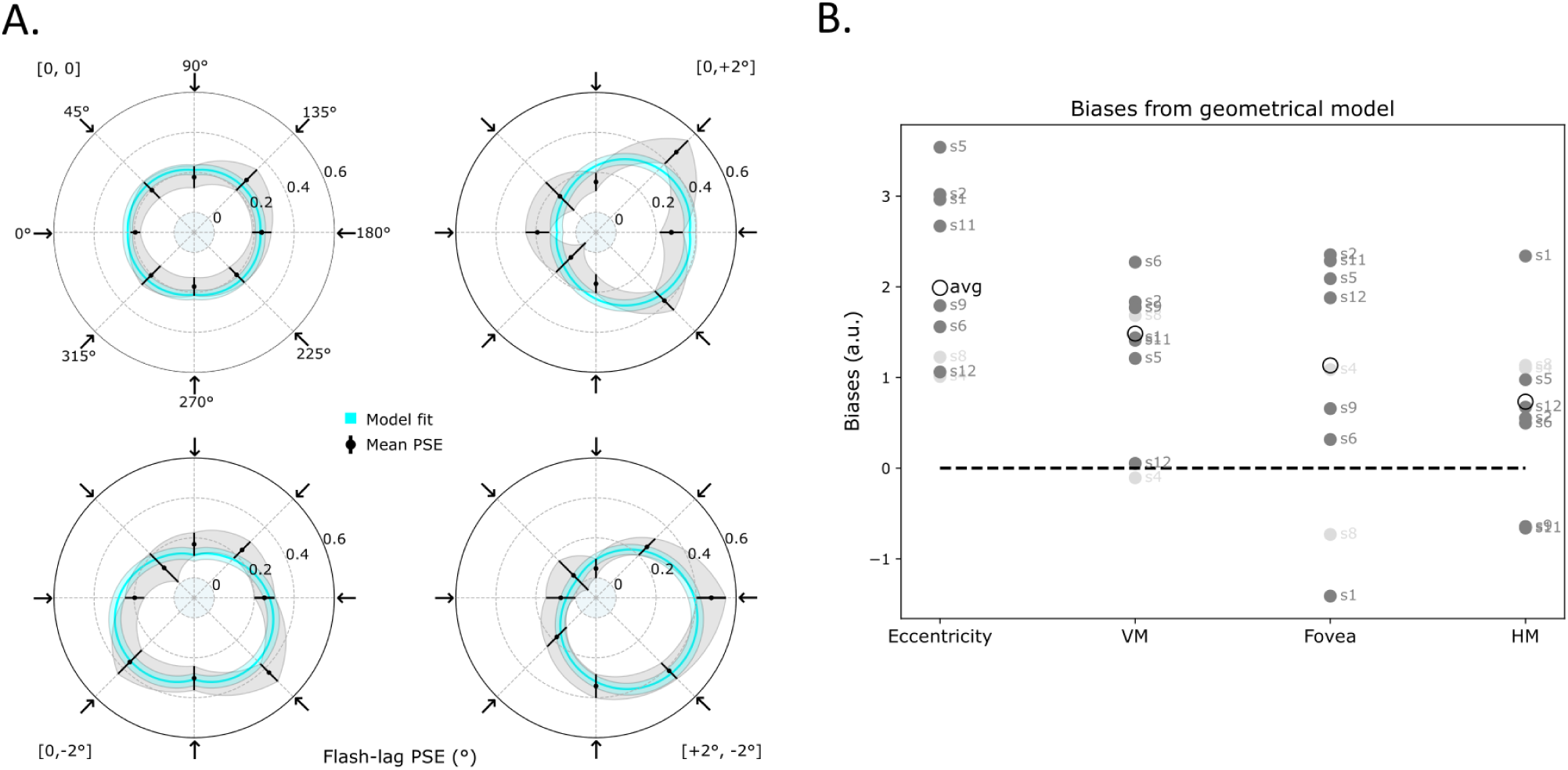
Experiment 4: Disentangling the effects of retinotopic features on FLE. **(A)** Polar representation of the strength of the flash-lag effect (FLE) in degrees (distance to center) as a function of the bar motion direction and value of bias of the geometrical model (angle of the plot). Each of the polar plots corresponds to one of the four positions where the FLE was measured: foveal [0°, 0°], on the VM and away from the fovea [0°, -2°], on the HM and away from the fovea [2°, 0°] or away from the HM, the VM and the fovea [2°, -2°]. The bar could move in eight different directions systematically shown by the arrows at the most external part of each polar plot (with the corresponding direction angle reported in the first upper left polar plot). Dots and error bars represent the mean ± standard error of PSE across subjects, the gray shaded area showing interpolated values between tested directions. The light-blue shaded disks in the center of polar plots represent FLE negative values (-0.1° to 0°). The light blue thick line represents the fitted model on the averaged data for each position, while the shaded area represents the 95% confidence interval of the model fit.. **(B)** The figure shows values of the four biases of the geometrical model: eccentricity, VM, fovea and HM. Dark grey dots represent individual subjects, labeled by subject number for which the fitting model was reported significant (with permutation test). Subjects 4 and 8 for which the fit does not perform better than for random sorted data are reported in light gray. The empty black dots show the averages computed across all subjects, used for the model fit representation of Panel A (the light blue thick lines).

VM-fugal and VM-petal motion (0° and 180° directions, respectively) at position [2°, 0] are not significantly different (0.19° vs. 0.28°, t(9) = 1.11, p = 0.30). Similarly, no significant difference was reported for VM-fugal motion (direction 0°) at position [2°, 0] and VM-petal motion (direction 0°) at position [0, 0] (0.19° vs. 0.19°, t(9) = 0.03, p = 0.98). However, the three conditions—VM-fugal and petal motion at position [2°, 0] (directions 0° and 180°, respectively) and VM-petal motion at position [0, 0] (direction 0°) in Experiment 2 (tested positions -2°, 0°, and +2°; see the first and last blue dots and the green dot in Figure 3)—lead to a significant difference, with a stronger FLE for HM. These discrepancies may reflect differences in the participant pool, as supported by the high inter-subject variability in Figure 5 for this condition, or in the specific grouping of conditions for the two experiments (see methods). Interestingly, our model predicts such a difference with a lower VM-fugal FLE than a VM-petal FLE (see the blue line at [0, 0], [2°, 0°] in Figure 5b).

The effect of moving towards or away from the HM is weaker compared to the other retinotopic features, with a stronger FLE for HM-petal conditions than HM-fugal conditions observed only for positions far from the HM: at [0°,-2°] (0.30° vs 0.17°, t(9)=3.45, p=0.009) and [+2°,-2°] (0.34° vs 0.05°, t(9)=3.45, p=0.002). The conditions +2° with downward motion (HM-fugal) and -2° with upward motion (HM-petal) of experiment 3, are similar to the 270° direction (HM-fugal) and 90° direction (HM-petal) at position [0°,-2°] of experiment 4. The trend observed in Experiment 3 between HM-petal and HM-fugal (0.26° vs 0.196°, p=0.23) became significant in Experiment 4 (0.30° vs 0.17°, t(9)=3.45, p=0.009).

### Geometrical modeling of physiological biases

All tested conditions of this experiment combine physiological biases related to three retinotopic features (fovea, VM and HM) and eccentricity. In order to disentangle, quantify and compare the respective effects on the FLE of visual motion toward (petal conditions) or away from (fugal conditions) these retinotopic features, we developed geometrical models of each of these biases and fitted their respective weights in the the global effect observed (see methods). In this model, we took into account the positive/negative variation of distance to these retinotopic features along the motion trajectories, assuming that the interaction between the three retinotopic features combined linearly. The fitted model (light blue curves in Figure 5) captured the overall trend of the group-averaged data (*R*^2^ = 0.768), with a permutation test confirming the statistical significance of the fit (p = 0, see method). We define the null hypothesis as the case where the data has no systematic relationship with the stimulus features, namely position and motion direction. Consequently, the model’s fit quality (R², rMSE) should be no better than expected from data in which these relationships have been randomly shuffled (see Methods for more details).

The permutation test was also applied at the single-subject level, rejecting the null hypothesis for 7 out of 9 subjects: only two subjects had p-values between 0.01 and 0.05 (sub12: *R*^2^= 0.234, p = 0.047; sub1: *R*^2^= 0.322, p = 0.015), while the remaining five had *R*^2^> 0.385 and p < 0.01. The model allowed quantifying for each individual subject as well as for the group average, how motion direction (towards or away from) relative to each retinotopic features (fovea, VM and HM) and how eccentricity can bias the FLE. Eccentricity gave the strongest bias on the group average (1.989), indicating that the FLE increases with eccentricity independently from its direction. For directional biases (towards or away from), by convention a positive bias indicates an increased FLE in petal vs fugal conditions. The strongest directional bias was observed for VM (1.484), then fovea (1.133) and the smallest was for HM (0.732). Analyzing these biases for each subject (Figure 5B), we found that VM and foveal biases are both stronger than HM biases in 6 subjects out of 9. Overall, there are no clear correlations between any of the biases. Data for subject 11 are shown in supplementary Figure 12, standard for average across subjects at supplementary Figure 13.

## Discussion

In this study, we sought to test the predictions of the PRM hypothesis according to which distinct processing for moving and static stimuli is due to the propagation of activity within retinotopic cortical maps. This creates an anticipatory wave in the direction of the moving object. According to this hypothesis, the discontinuities and deformation of cortical retinotopic maps should have a measurable impact on the FLE, the first leading to a drastic reduction of the effect and the latter to a modulation. We designed psychophysical experiments to precisely test these predictions on humans.

We report a drastic drop in the FLE when the moving stimulus crossed the VM. This effect occurred regardless of motion direction (i.e. the same for rightward and leftward motion) and is also independent of trajectory length. Then we systematically evaluated the FLE for horizontal and vertical motion, both with tested positions before and after crossing the VM and the HM. The drop in FLE observed for horizontal motion when crossing the VM in Experiment 2 is very fast and occurs within less than 1° of visual angle. Notably, the effect does not fully return to pre-crossing values: the FLE remains consistently lower when moving away from the VM than when moving towards VM. Conversely, for a bar moving vertically, the FLE increases when the bar crosses the HM for both upward and downward motion when measured at low eccentricity (0.5°). In this region of the visual field we showed that the FLE is also higher for downward than for upward motion. In these two later experiments, the fovea was also crossed together with meridians. In order to tease apart various components that can influence the FLE, we designed an experiment to quantify the respective effects of eccentricity, motion direction approaching or receding either of the three retinotopic features (fovea, VM and HM). Overall, we report an increase of the FLE with eccentricity and for directions towards (petal) visual features compared to away from (fugal) directions. Ultimately, we modeled the behavioral data and found that the FLE depends on a variety of uncorrelated biases, primarily related to eccentricity and VM with foveal biases and HM biases being less important. This is the first study that investigates how the FLE is modulated by crossing and the directionality of motion to retinotopic features, and how the effect evolves with eccentricity. Crucially, the originality of our approach lies in the usage of a Bayesian adaptive method (see Methods), that enabled fine-grained measurement of the FLE across many positions in the ±2° region around fixation. This provided unprecedented resolution for characterizing FLE evolution in central vision.

Although not investigating the FLE per se, some studies have probed the effect of crossing retinotopic features on other visual illusions. Liu, Tse, and Cavanagh (2018) used a double-drifting gabor moving in the two cardinal and two obliques directions, for investigating the role of VM and HM on the global motion perception. To avoid any effect of the movement direction with respect to the fovea, the authors used motion trajectories back and forth. Liu et al. showed that the typical illusory-position shift of the stimulus is reduced when the illusory motion does not cross the VM and HM. They attributed such an effect to the interareal connectivity discontinuities at the level of HM for the visual area V2 and V3, and the contralateral lateralization of the visual areas mediated by the corpus callosum, with a representational discontinuity corresponding to the VM. Our study is, to our knowledge, the first to measure the FLE explicitly when crossing the fovea, VM, and HM. In this context, only few studies have focused on vertical motion and crossing of HM (Nagai et al. 2002; Ichikawa and Masakura 2006; 2010). Among these, Nagai et al. tested vertical motion at three different positions –pre-crossing, foveal, and post-crossing– along the VM at 0° eccentricity, but at higher elevations (±3.05°). While the eccentricity range matches that of our study, the authors pooled data across positions, making it impossible to isolate effects at specific test positions. Ichikawa et al. instead presented the stimuli both on the right and left of the VM, at eccentricities of ±2.5°, beyond the range investigated here.

The role of motion direction to retinotopic features in modulating the FLE has been investigated, particularly for horizontal motion. Consistent with previous studies (Mateeff and Hohnsbein 1988; Kanai et al. 2004; Shi and Nijhawan 2008), we observed a generally stronger FLE for horizontal motion directed toward the VM (i.e., VM-petal), compared to motion directed away (VM-fugal), as shown in Experiments 1, 2 and 4. Our findings are in line with those of Shi and Nijhawan (2008), who employed a similar stimulus configuration and observed comparable FLE asymmetries. Interestingly, these authors reported negative FLE values for foveo-fugal motion, particularly in the right hemifield, a pattern that resonates with the variability observed across subjects in our VM-foveo-fugal condition (Experiment 2, Figure 2, green dot). In contrast, the influence of motion direction relative to the HM (i.e., vertical motion) has received less attention. Most studies focusing on vertical motion have examined absolute direction effects (upward vs. downward) rather than HM-petal/fugal motion. For instance, Nagai, Kazai, and Yagi 2002 reported a larger FLE value for downward motion than upward. This directional asymmetry might reflect higher-level processes influenced by a prior knowledge of physical gravity –i.e. representational gravity, for review see Hubbard 2020–. Supporting this hypothesis, Nagai et al. demonstrated that the downward motion-direction effect is positively modulated when aligned with the environmental vertical but not with the body vertical axis. On the other hand, Ichikawa and Masakura 2010 did not observe any significant difference between upward and downward motion. These conflicting findings may be reconciled by our observation that directional asymmetries in vertical motion were evident within a narrow range around the fovea and HM (±0.5°), but not outside of this region, suggesting a highly localized effect that previous studies may have missed.

Unlike previous reports, we did not observe significant difference in FLE between the left and right visual fields (Figure 2B and supplementary Figure 1). This contrasts with several studies that have reported a larger FLE in the left visual field compared to the right (Kanai et al. 2004; Shi and Nijhawan 2008; Suzuki et al. 2023). A likely explanation lies in eccentricity differences: most previous work tested at larger eccentricities (at vertical and horizontal eccentricities >2°), while our study examined FLE in a narrow region around the fixation point, with stimuli positioned within ±2° of the VM and HM. This interpretation is further supported by findings from Amadeo et al. (2022), who similarly tested within a central range (1.1° to 2.5° from fixation) and did not report significant effects of motion directions across different hemifields, suggesting that hemifield asymmetries may be diminished or absent at low eccentricities.

We also examined how the FLE varies with eccentricity. To the best of our knowledge, only a few studies have focused on this feature (Baldo et al. 2002; Linares et al. 2007) focusing mostly on the relative eccentricity between target and flash stimulus, not to their absolute position in the field of view. In particular, Baldo et al. 2002 attributed the increase of FLE to a low position predictability of the flashed object (and conversely, a decrease in FLE might be due to a larger position predictability of the flashed object). Further, only a few studies have specifically investigated the FLE at central visual field locations, and most with paradigms not directly comparable to ours. For instance, although Kanai, Sheth, and Shimojo 2004 used a design with stimuli at 0° elevation and horizontal motion, the FLE was evaluated in a flash-terminated task, while Shi and Nijhawan 2012 explored the foveal region using a small dot stimulus (rather than a bar), presented between 0° and 2.3°, in flash-terminated and flash-initiated tasks. Among studies with more comparable configurations, Amadeo et al. (2022) tested both a peripheral and a central condition; in the latter, stimuli were presented between 1.1° and 2.5° of eccentricity with a moving bar positioned 2° below fixation and a reference target appearing at the midpoint of the trajectory. They reported a significant difference in FLE between peripheral and central conditions, for both the tested subject groups (early deaf and hearing participants), with higher FLE for peripheral than central, both in VM-petal direction. Their results are in line with those observed in our study, further supporting the reliability of our findings at low eccentricities.

### Possible underlying mechanisms

The remainder of this discussion will focus on identifying the possible mechanisms underlying the effects reported in this project. The main PRM hypothesis guiding this study posits that the FLE originates from the propagation of neural activity within retinotopic maps. The PRM hypothesis predicts that the FLE should strongly depend on the topology of retinotopic maps, canceled by retinotopic discontinuities and modulated by retinotopic anisotropies. We distinguish three overlapping, yet distinct, factors: crossing a retinotopic feature, eccentricity, and motion direction relative to these features (petal vs. fugal). While often entangled, each of these factors can affect the FLE through distinct mechanisms.

Crossing a retinotopic feature implies a transition between functionally distinct cortical territories, which may also be anatomically distinct. This is particularly evident at the vertical meridian (VM), where interhemispheric communication is required. The VM separates the left and right visual fields, reflecting an anatomical split of the two hemifields, each represented in contralateral cortical visual areas (Hubel and Wiesel 1967; Innocenti et al. 2022). Part of V1at the boundary with V2 –corresponding to the retinotopic location of the VM in early visual areas– sends callosal projections to the contralateral hemisphere, specifically at the homologous area V1-V2 border. Hence the VM is represented in both hemispheres, extending slightly into the ipsilateral side, within a narrow band of ±1° around the VM in human and NHP (Essen et al. 1982; Kennedy et al. 1986; Kennedy and Dehay 1988; Clarke and Miklossy 1990; Zilles and Clarke 1997) . It is therefore crucial to understand the distinct properties of the two major types of cortico-cortical connections involved: intra-cortical horizontal and inter-cortical callosal. The former span approximately 5-6 mm and conduct signals at speeds around 0.3-0.5 m/s in NHP (Grinvald et al. 1994; Angelucci et al. 2002; Reynaud et al. 2012; Muller et al. 2014) . In contrast, callosal fibers cross the midline, are roughly ten times longer and myelinated so propagate at higher conduction velocities ranging from 4.9 to 11.4 m/s in NHP (Caminiti et al. 2009; Tomasi, Caminiti, and Innocenti 2012; Caminiti et al. 2013). Therefore, when a moving stimulus crosses the VM, visual information near the VM (±1°) will not propagate through horizontal axons anymore but through callosal pathways. Benvenuti et al. (2020) found in NHP V1 that crossing the VM disrupts the observed latency reductions along trajectories restricted to the contralateral hemifield, producing no changes of the neuronal response. These findings support the hypothesis that horizontal connectivity plays a central role in modulating neuronal activity, potentially at the origin of the latencies reported in the FLE. The drop in FLE we observed when a moving stimulus crosses the VM is comparable with the observations reported in Benvenuti et al (2020). The transition to callosal connectivity at the VM seems to interrupt the latency-based accumulation process that could be at the origin of the illusion, resulting in a sudden change in latency decrease and, consequently, a reduction in the perceived spatial shift.

The horizontal meridian (HM) marks the boundary between the upper and lower visual fields, but unlike the VM, it corresponds to an anatomical division only for areas downstream of V1. Anatomically, the HM separates upper and lower visual field representations across ventral and dorsal portions of contralateral visual areas respectively. In area V1, both fields are represented in a continuous map along the calcarine sulcus. Here, contrary to what was observed for the VM crossing, we observed an increase in FLE when stimuli crossed the HM. To the best of our knowledge, no studies have systematically examined cortical responses to moving stimuli at HM crossings, either in early or higher visual areas. One possibility is that this increase in FLE at the level of the HM may reflect transitions between dorsal and ventral processing streams, suggesting the involvement of higher-level visual areas and their corresponding retinotopic representations. However, why this results in increased FLE remains unclear and calls for further investigations.

Another effect we wished to probe in the PRM hypothesis, is the effect of cortical magnification factor, which reflects the disproportionate allocation of cortical surface to central compared to peripheral visual space. The magnification factor decreases exponentially along eccentricity for both NHP and humans, although with higher values in humans (Van Essen et al. 1984; Harvey and Dumoulin 2011). The horizontal connections in V1 span across a cortical distance (i.e.∼5-6 mm) and speed (between 0,3-0,5 m/s) that are, in a first approximation, identical across the V1 retinotopic map. Since cortical magnification decreases with eccentricity, these connections cover a larger portion of equivalent visual space in the periphery than in the fovea. Consequently, the corresponding visual propagating waves should become faster and extend farther for more eccentric retinotopic positions. Close to the fovea, the limited extent of propagation in the visual space and its smaller equivalent visual speed should result in weaker anticipatory effects. This matches our results, captured in the last experiment which clearly shows an inverse dependence of the FLE with eccentricity. It may also contribute to explain the difference of FLE effect in experiment 3 with the upper visual field yielding higher FLE than the lower visual field. Indeed, it has been shown that a larger cortical surface in V1 is allocated to the lower rather than the upper VM (Himmelberg, Tünçok, et al. 2023). However, there is potentially another confound in comparing the lower and upper hemifields in Exp 3: the bar was moving downwards in the upper hemifield and upwards in the lower hemifield. It could well be that there is another factor explaining the difference of FLE in these two conditions, like a “gravity” bias, known in literature as representational gravity (Hubbard 2020). This has already been reported in FLE only at certain specific conditions (Nagai et al. 2002).

We observed a systematic effect of motion direction relative to retinotopic features (VM, Fovea and HM), the FLE being larger when moving towards these than away. Since the trajectory activates the same retinotopic region, the main difference lies in the order of the sequential activation along that cortical territory. As the magnification factor increases closer to these retinotopic features, the sequence of cortical activation accelerates when it approaches them and slows down when it moves away from them. Since the FLE is larger when it moves towards, one hypothesis is that the beginning of the trajectory, which will move faster and further away in the cortical space, may play a more important role in influencing the FLE than the end of the trajectory. Another, complementary, mechanism could arise from callosal modulations of stimuli that start from the vertical meridian. Studies in cats and monkeys indeed show that callosal deactivation affects responses to VM-fugal but not VM-petal visual motions (Conde-Ocazionez et al. 2018; Raemaekers et al. 2009). How this modulation could affect the FLE remains to be understood, but this result shows a differential modulation of the response of VM-fugal and VM-petal stimuli.

Using psychophysics, we have verified the two predictions that follow from the PRM hypothesis, namely a cancellation of the FLE when crossing a visual feature that corresponds to retinotopic disruption, and modulations of the FLE by retinotopic anisotropies. However, the specificities of the effect, with a strong reduction induced by the VM crossing and a lack of reduction induced by the HM crossing, suggests that the FLE effect is driven by propagation of activity within V1 retinotopic maps, the only area that has a disruption along the VM and not the HM. The clear eccentricity effect of the FLE also pleads in favor of early visual cortices since the retinotopic representation in higher order areas can follow a more complex pattern than the one observed in V1, although with a similar magnification factor decrease with eccentricity (Harvey and Dumoulin 2011). In the NHP, complex retinotopic representations have been reported for visual areas downstream to V1, starting from V2, which is known to have a patchy and sinusoidal organization in the tree-shew (Sedigh-Sarvestani and Fitzpatrick 2022) or dorsal visual areas like V3 in primates (Shipp et al. 1995; Yu et al. 2019; Ribeiro et al. 2023) or even higher level areas (V4, MT, etc.), could have some effects on the seen effect. To fully understand this phenomenon, further investigation into the unique retinotopic mapping properties of each visual area is required that could help drawing more precise predictions under the PRM hypothesis.

## Conclusion

In conclusion, our findings support the PRM hypothesis and suggest that horizontal intra-cortical connectivity, particularly within V1, plays a central role in the emergence of the FLE. Using a Bayesian adaptive method and hypotheses derived from mesoscopic NHP imaging, we demonstrate that fine-scale cortical structure can shape perception at a local level. Notably, this shows that the FLE strength is not uniform in central vision but varies sharply depending on proximity to and crossing of retinotopic features. This calls for a more comprehensive investigation of how psychophysical measures may be affected by discontinuities in cortical maps. Moreover, individual variability, such as the ones we observed here for the strength of the FLE, may reflect anatomical individual differences. Further work, combining non-invasive neuroimaging with psychophysics could further uncover how cortical idiosyncrasies shape perception. We suggest that this approach, testing the inferences of how cortical maps topology can influence our perception, could help to better understand the cortical origin of psychophysical observations.

## Material and Methods

### Ethics Statement

For each experiment of this project, participants gave a prior written consent after being informed of the methods used and their right to interrupt if they wished. Participants were compensated for each experimental protocol. This project was approved by the Comité d’éthique d’Aix-Marseille Université (reference 2014-12-3-06) and complies with the regulations described in the Declaration of Helsinki (2012).

### Subjects

We collected data from a total of 24 subjects. Of these, only two subjects participated in all four experiments, both of whom are among the authors. We analyzed data exclusively from subjects for whom the Bayesian adaptive method successfully converged (see the Bayesian Adaptive Method section). In Experiment 1, data from eleven subjects were collected; however, only ten participants’ data were analyzed, with a female ratio of 8/10. The mean age of the subject pool was 31.1 years (range: 24–40). In Experiments 2 and 3, data from the same twelve subjects were collected, but only ten participants’ data were analyzed, with a female ratio of 6/10. The mean age of the subject pool was 27.4 years (range: 21–41). Nearly all participants completed both experiments on the same day, except for one who was tested two weeks apart. Experiment 4 required a larger pool of participants: out of 16 subjects, data from 9 subjects were retained, with a female ratio of 4/9. The mean age of the subject pool was 29.8 years (range: 21–41). Due to the large number of conditions, the testing was distributed across four different sessions. The mean time difference between the first and last session was 59.6 hours, with a maximum of 140 hours and a minimum of 24 hours. All participants had normal or corrected-to-normal vision (contact lenses were allowed, but glasses were not).

### Apparatus

Participants sat in a dark room in front of a screen, with their head movements restricted by a chin and forehead support, placing their eyes at 57cm from the screen center. Stimuli were generated on a computer running Windows 7 operating system, with routines written in Python and using the core libraries of PsychoPy (Peirce 2007). Visual stimuli were displayed on a Samsung TFT monitor running at a resolution of 1680×1050 and refreshed at 100Hz (frames of 10 ms), over a uniform 50% grey background (25.8 cd/m2 luminance). The right eye position was monitored by a Tobii X120 video eye tracker (sampling at 120Hz), During trials, participants were asked to look at a central fixation point represented by a small white disk (0.2° in diameter). Trials were aborted and repeated later (two trials ahead when possible) when fixation was broken, that is, when horizontal or vertical error was greater than 0.5° and 1.2°, respectively, for at least 42ms (5 tracking frames).

### Procedure

#### General Visual Stimulation and Task

We conducted four experiments to investigate specific aspects of the Flash Lag Effect (FLE). The visual stimulation always included a white fixation point on a grey background; a white bar moving along a linear trajectory at a constant speed of 6.6°/s; and a single or a pair of squares flashed for 10ms (one frame) (Figure 2A schematic of visual stimulation). The relative position of the flash and the bar was varied using a Bayesian Adaptive Method (see below). For Experiments 2, 3 and 4, a pair of squares were flashed on a virtual line parallel to the bar, above and below its trajectory. For Experiment 1 only one square was flashed above the bar. In all experiments the flash was presented at the midpoint of the bar trajectories and the task was to compare in a 2AFC the bar position relative to the flash in space. They were asked: “At the moment the flash occurred, where was the bar relative to the flash?” Responses were given either with the keyboard arrows or the numerical keypad depending on the experiment. At the beginning and at the end of the trajectory we implemented a bar fading (in and out) duration of 100ms each for experiment 2, 3 and 4. To prevent cognitive strategies while allowing sufficient response time, the reaction time (RT) was limited to a maximum of 2.5 seconds but with a general lower RT across participants and experiments (see supplementary). The bar disappeared either after the participant gave their answer or after having traveled a maximum of 10° (2.5s after flashes). Trials started automatically 2 seconds after the previous answer or maximum reaction time was reached. Trials were grouped in blocks of a subset of conditions so that the participants could focus their attention to a limited area of the visual field. At the start of each block, an instruction was displayed to indicate the tested spatial area. The participant could take a short break between blocks

#### Bayesian Adaptive Method

For each tested condition, an optimal Bayesian adaptive method was used to determine the tested positions of the flash relative to the bar (Kontsevich and Tyler 1999). After each trial is answered, the method provides an estimation of both the point of subjective equality (PSE) and the standard deviation (SD) of the psychometric curve (a cumulative Gaussian), together with their respective theoretical 95% confidence intervals. Based on these, it computes the best value in the test range to be used in the next trial to maximize the gain of knowledge. This fast-converging method sampled 100 values for the flash position in the range from -1.982° and +1.32° (corresponding to a flash position that ranged from a -300ms lag to a +200ms lead).

#### Experiment 1: Horizontal Motion and Flash Timing Relative to the Vertical Meridian (VM)

In Experiment 1, we tested the effect of a bar moving rightward, at the crossing or not of the VM (Figure 2). The traveled distance before the reference flash occurred could be 1°, 2°, 3° and 4°. The flash appeared either 0.5° before or after the bar crossed the hemifield boundary (the VM), resulting in eight conditions: four with the bar crossing the VM and four without. The bar disappeared either after the participant answered or after traveling an additional distance of 1.5°. The bar was centered at -4° of elevation with a height of 4° and width of 0.5°. The single square flashed appeared above the bar at variable offsets (side of 0.5° and separated from the bar by 0.75°). Trials were blocked in groups of 64 in which the tested spatial position was always the same, either -0.5° or +0.5° regardless of the travel distance. Blocks were alternating with the conditions of the initial block balanced across participants.

#### Experiment 2: Flash Position and Initial Eccentricities Relative to the VM

Building on Experiment 1, in Experiment 2 the flash-lag effect was measured after a fixed travel distance of 2°, but at varying eccentricities ranging from -2° to +2° in steps of 0.5° to allow for a fine characterization of the effect reduction observed when crossing the VM. This setup resulted in eight conditions, corresponding to the eight tested positions. The bar (3° in height, 0.5° in width) traveled rightward along the HM. Two squares (side of 0.5°) were flashed at the moment when the bar had traveled 2°, above and below the bar (separated by a 0.5° gap) with a variable offset. Trials were grouped in blocks of 72 in which the tested spatial positions were always in the same area, either to the left (eccentricities -2° to -1°), or to the center (-0.5° to +0.5°) or to the right (+1° to +2°). The order of the first three blocks was randomized across participants and repeated for the remaining blocks of the experiment.

#### Experiment 3: Vertical Motion and Flash Timing Relative to the Horizontal Meridian (HM)

Similarly to Experiment 2, in Experiment 3 the flash-lag effect was measured after a fixed travel distance of 2°, at varying eccentricities (-2°, -0.5°, +0.5° and +2°) to allow a characterization of possible effects given by crossing of the HM. The bar (3° in height, 0.5° in width) traveled along the VM, both in upward and downward directions. Similarly to the previous experiment, this setup resulted in eight conditions: four tested positions and two motion directions. Two squares (side of 0.5°) were flashed at the moment when the bar had traveled 2°, to the right and left of the bar (separated by a 0.5° gap) with a variable offset. Trials were grouped in blocks of 64 in which the tested spatial positions were always in the same area, either above (eccentricities of -2°, -0.5°) or below (+0.5° and +2°) regardless of motion direction. Blocks were alternating with the conditions of the initial block balanced across participants.

#### Experiment 4: Flash Position Proximity to Fovea, VM, and HM

To investigate the effects of proximity to the fovea, VM, and HM, and potential interactions between these factors, we designed Experiment 4. In this experiment, the moving bar could have one of eight directions (from 0° to 360° in steps of 45°). The bar (1.8° in height, 0.3° in width) moved towards four different tested positions: on the fovea ([0°, 0°]), on the VM ([0°, -2°]), on the HM ([2°, 0°]) and away from fovea, VM and HM ([2°, -2°]). After traveling a visual distance of 2°, the bar reached the tested position and two squares were flashed (side of 0.3°) at each ending of the bar (separated by a 0.5° gap) with a variable offset. The experiment has a total amount of 32 conditions. Trials were grouped in blocks of 32 in which the tested spatial position was always the same (e.g. [0, 0], [+2, 0], [0, -2] or [+2, -2]) so that attention was spatially focused on the same location, and the motion direction was either cardinal (0°, ±90°, and 180°) or oblique (±45° or ±135°). Answers were given using the numerical keypad of the keyboard, and participants were instructed to keep four fingers on the corresponding directions (8/2 and 4/6 for cardinal or 1/9 and 7/3 for oblique motion directions, respectively). At the start of each block, an instruction was displayed to the participant with a scheme indicating the tested spatial location, the 4 possible motion directions and its corresponding keys. In each block, conditions were repeated 8 times. The order of blocks was determined so that the two motion directions (cardinal and oblique) in a given spatial location were tested successively, and the spatial location tested was assigned to subject numbers using a 4×4 latin square design to cancel for possible order biases in the results. Each experimental session contained 24 blocks, for a subtotal of 768 trials. The experiment contained 4 sessions, for a total of 3072 trials including 96 repetitions of each condition.

### Data analysis

#### Non-linear Regressors Model

To analyze the responses to a moving bar stimulus across different positions in the visual field (as described for the experiment 4), we developed a set of parametric models based on the spatial configuration and movement directions.

For each of the four conditions (foveal, centered on vertical meridian, centered on horizontal meridian, away from fovea, horizontal and vertical meridian), we modeled the perceptual responses as a function of the stimulus direction using nonlinear regression functions (see Supplementary 11C). The general form of the models incorporates a combination of cosine and sine components, capturing the horizontal and vertical motion contributions. The principal rationale is to compute the *variation of* trajectory to a retinotopic feature (being the two cardinal meridians (horizontal *H* and vertical *V*) and the fovea-fixation point (*F*)). These components are equally weighted by the eccentricity of the trajectory, represented by an absolute distance between the fixation point and the mean point of the bar trajectory: the eccentricity is indicated as *ecc*, with trajectory length indicated by *a* (see Supplementary 11B).

We formalize the model as:

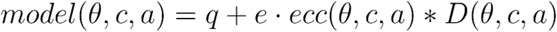

With as baseline *D(θ, c, a)* and distances from horizontal, vertical meridian and from fovea:

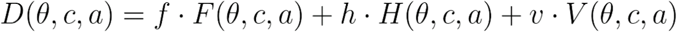

With *q* as baseline and *D(θ, c, a)* istances from horizontal, vertical meridian and from fovea:

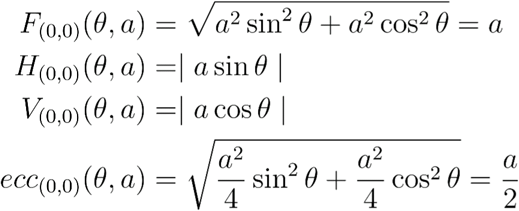

The independent variable θ corresponds to the motion direction. It corresponds to the 8 directions between 0° and 360°, stepping 45°. *c* is the center of stimulus presentation (*x*_0_, *y*_0_): it corresponds to the position *when* the flashing reference occurs and we impose that at this point the bar has traveled for a length *a*. The model finds the set of values *q, e, f, h, v* that better approximate the results obtained from subjects in the experiment 4. Below the components *F(θ, c, a), H(θ, c, a), V (θ, c, a)* are described. For the 4 positions employed in the experiment 4, we have for ***foveal*** ergo *c* = (0):

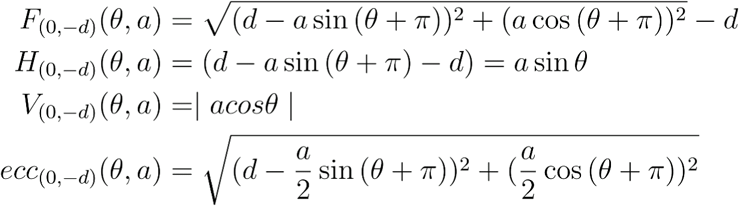

For ***centered on horizontal meridian***, ergo *c* =(*d*, 0):

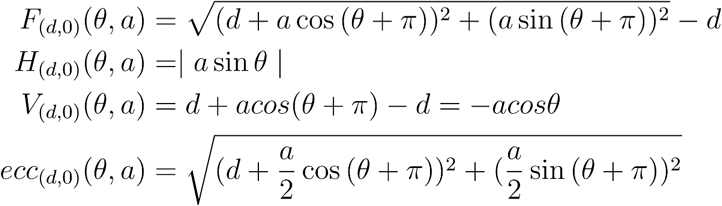

For the case ***away from fovea, horizontal and vertical meridian***, ergo *c* =(*d*, – *d*):

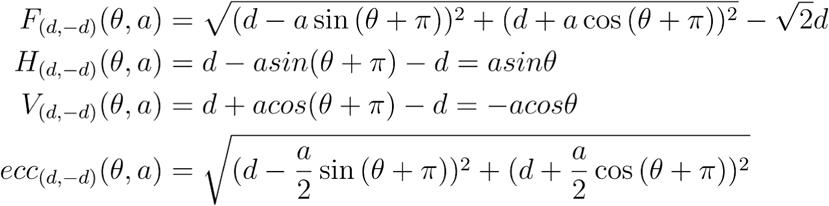

For all the non foveal positions a component π is added to the θs, changing the sign of the equations: this is due to the assumption that the biggest FLE is associated with foveo-petal directions.

The margin of error of the fit is computed as the half-width of the confidence interval (the light blue shaded area in Figure 5A), with a 95% of confidence level.

The code for fitting analysis and mathematical implementation of the equations is available at the repo https://github.com/SaGiancani/model-fle

#### Permutation test on model fitting

We systematically shuffled the data’s relationship to the stimulus features (random attribution of position and direction) to assess the model’s statistical significance by comparing the fit quality – fit r-squared (*R*²) and rooted mean squared error (rMSE)– for shuffled data and actual data. Data points from the psychophysical data of Experiment 4, at a single subject level and on the group average, were shuffled across motion direction and position and then fitted according to our geometrical model. The process was repeated 1,500 times. P-values were calculated for each subject and for the overall average across subject distributions to evaluate statistical significance. Code and results for the permutation analysis of model fitting are available at https://github.com/SaGiancani/model-fle.

## Acknowledgment

The authors would like to thank Martin Szinte for fruitful exchanges for the drafting of the Discussion section. This project has received funding from the European Union’s Horizon 2020 research and innovation programme under the Marie Skłodowska-Curie grant agreement N° 956669, as well as a grant by Fondation Berthe Fouassier/Fondation de France to S.G. and F.C..

All authors have approved the final manuscript and declare no competing interests. The work is original and has not been published elsewhere, nor is it currently under consideration for publication.

## Supplementary figures

**Supplementary. Figure 1.**
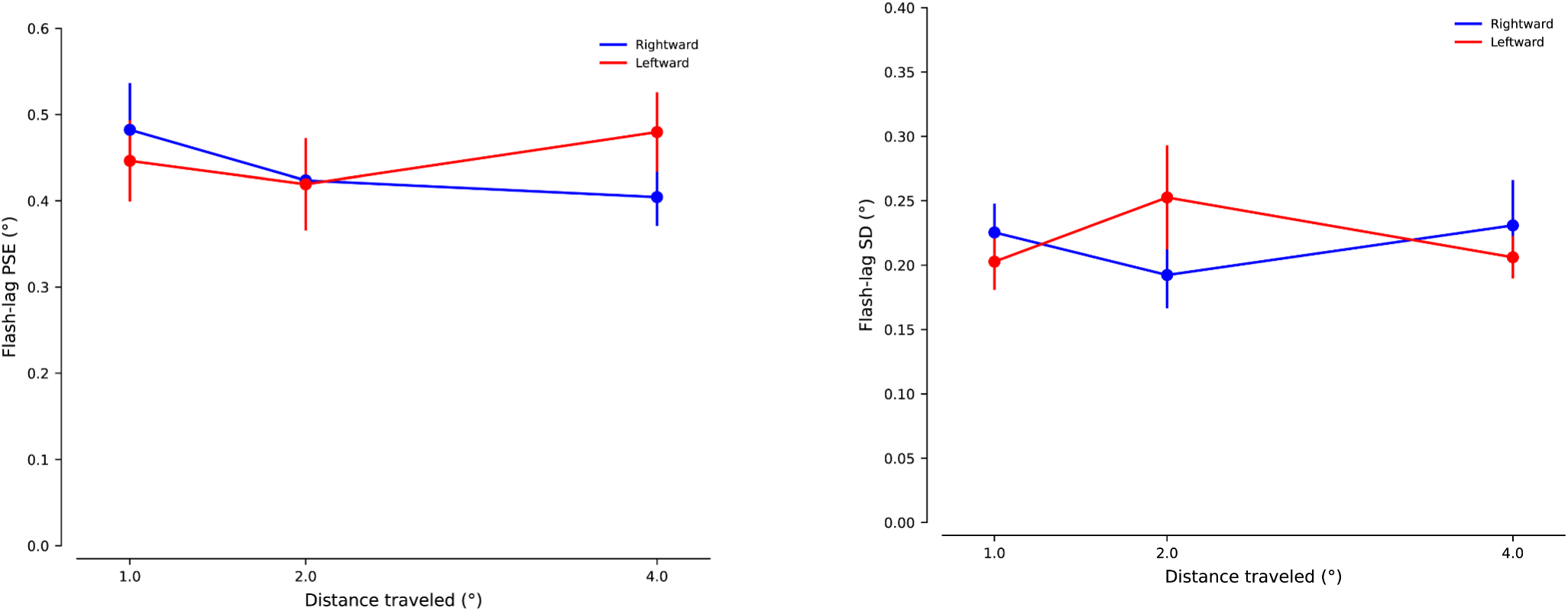
Control experiment comparing: leftward motion in the right hemifield (red) vs rightward motion in the left hemifield (blue). Average (left) and standard deviation across subjects (right) (in °).

**Supplementary. Figure 2.**
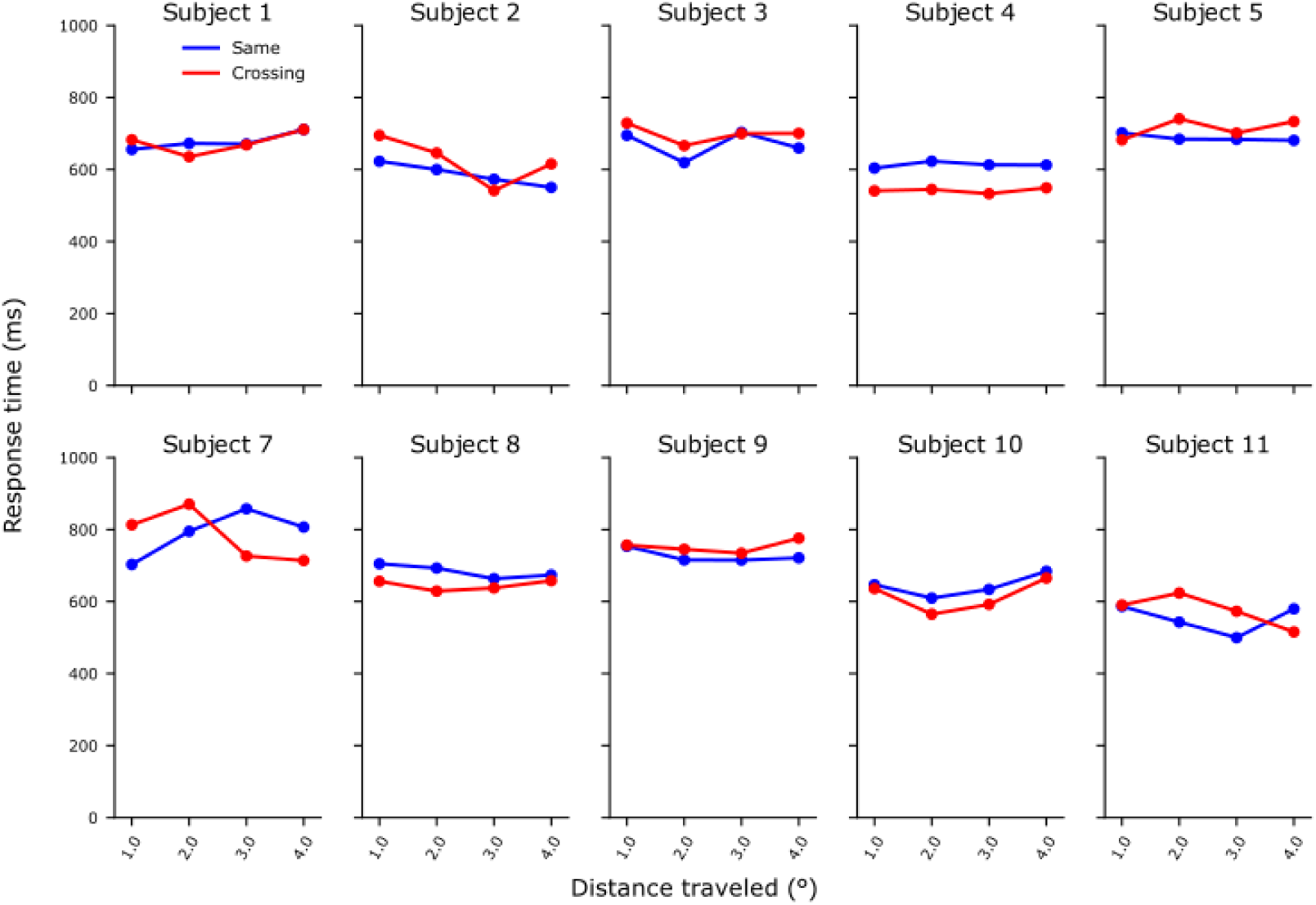
Experiment 1: Flash-lag response time individual results.

**Supplementary. Figure 3.**
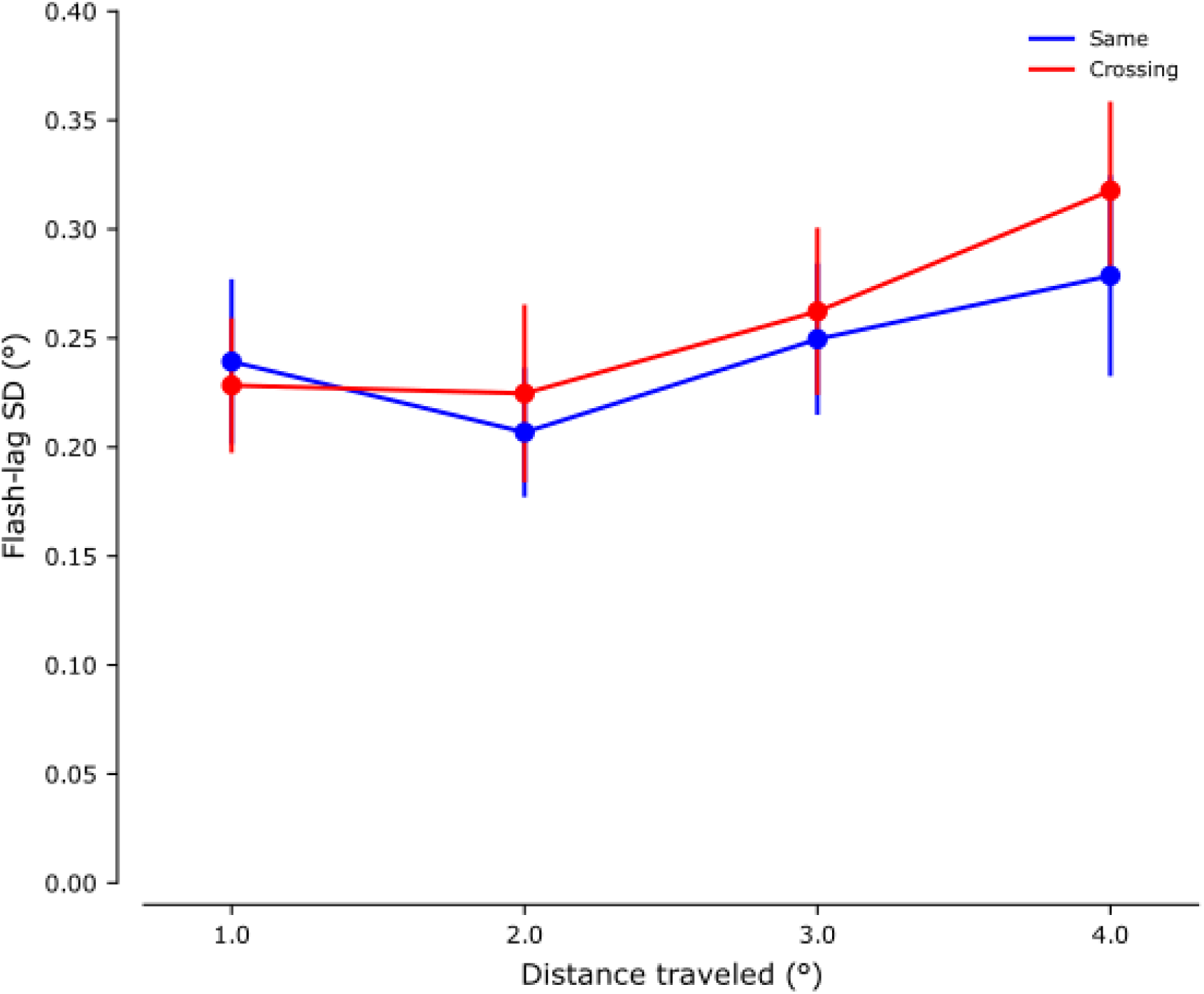
Experiment 1: Average flash-lag standard deviation (in °).

**Supplementary. Figure 4.**
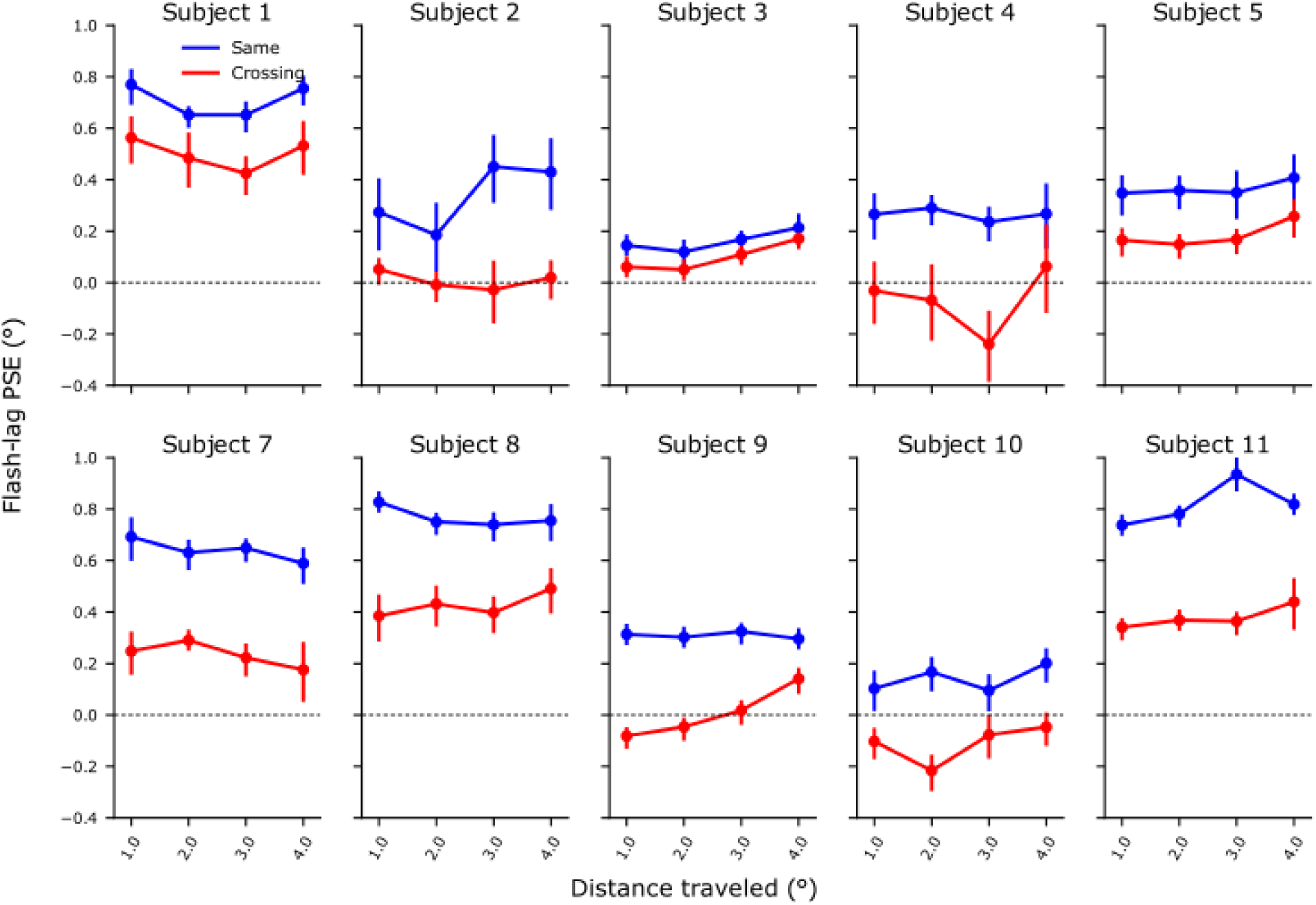
Experiment 1: Flash-lag PSE individual results (in °).

**Supplementary. Figure 5.**
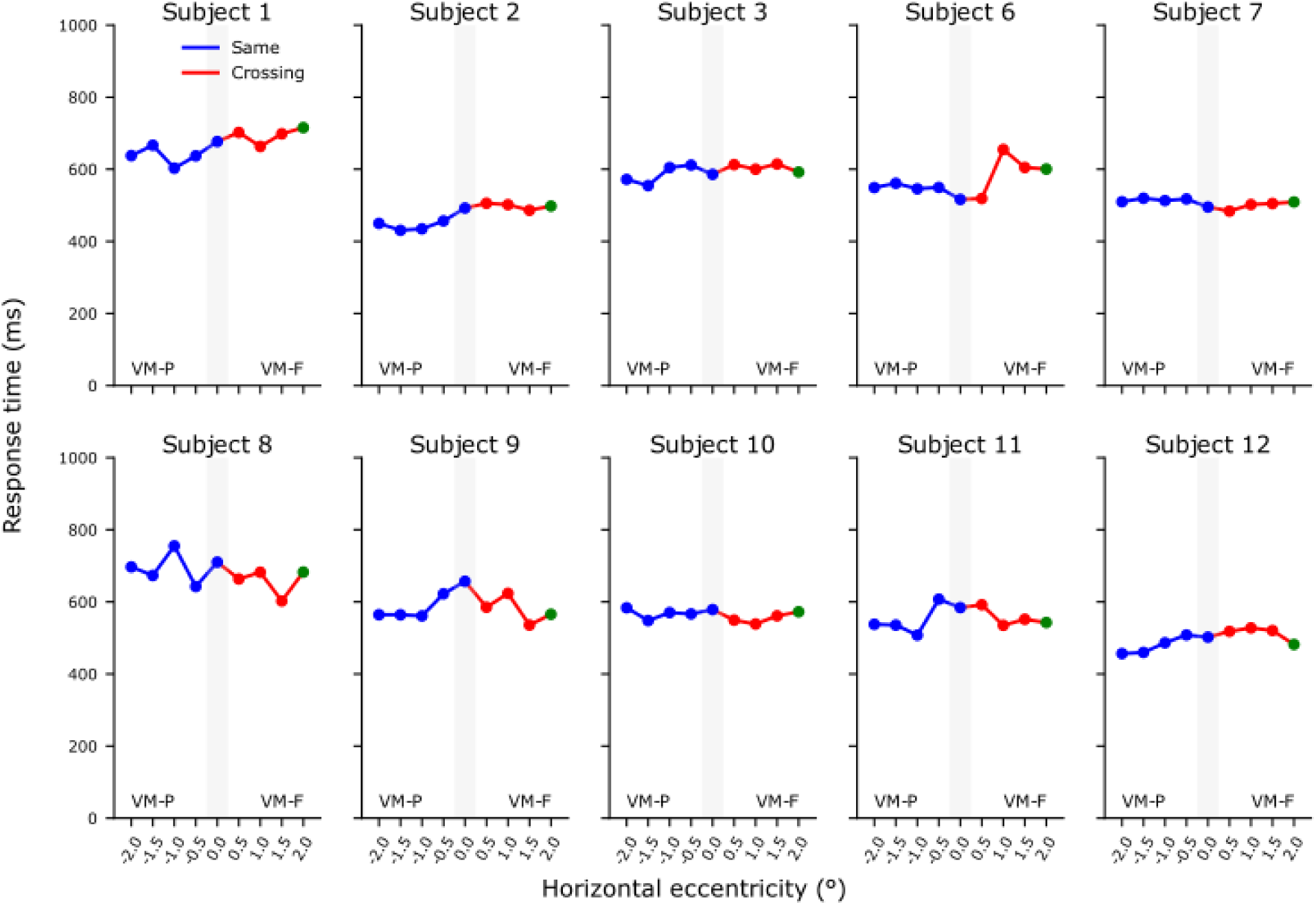
Experiment 2: Flash-lag response time individual results (in ms).

**Supplementary. Figure 6.**
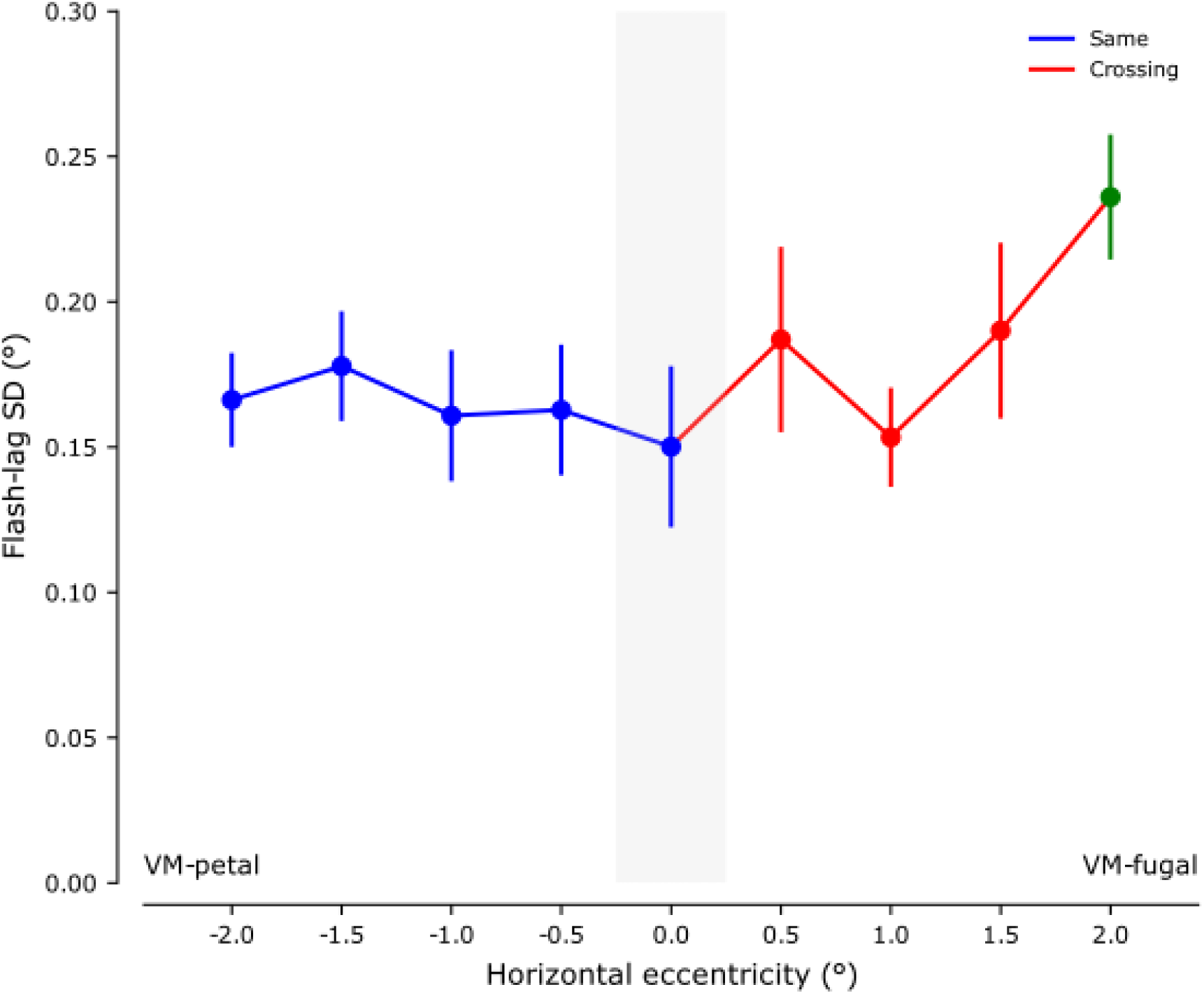
Experiment 2: Average flash-lag standard deviation (in °).

**Supplementary. Figure 7.**
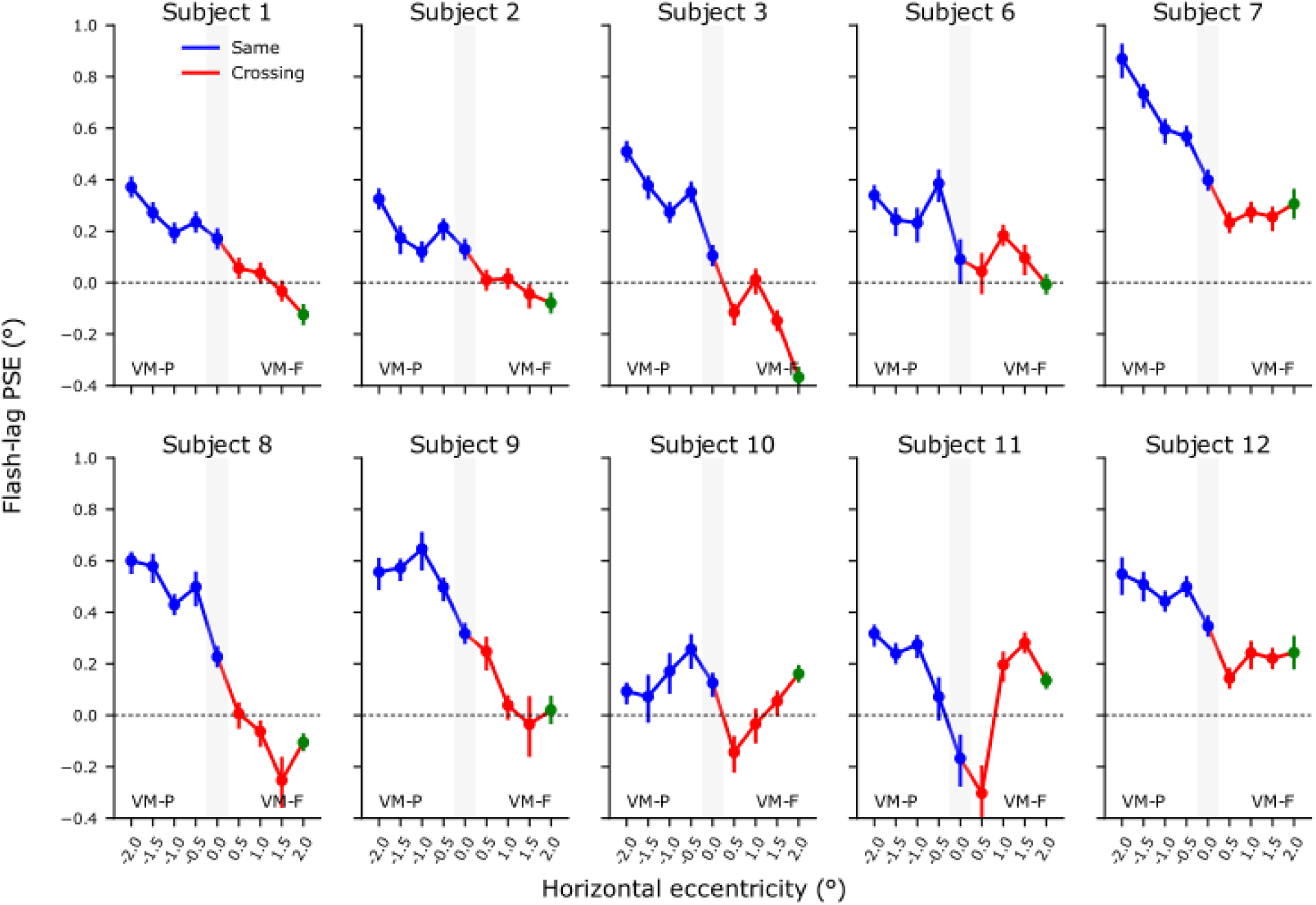
Experiment 2: Flash-lag PSE individual results (in °).

**Supplementary. Figure 8.**
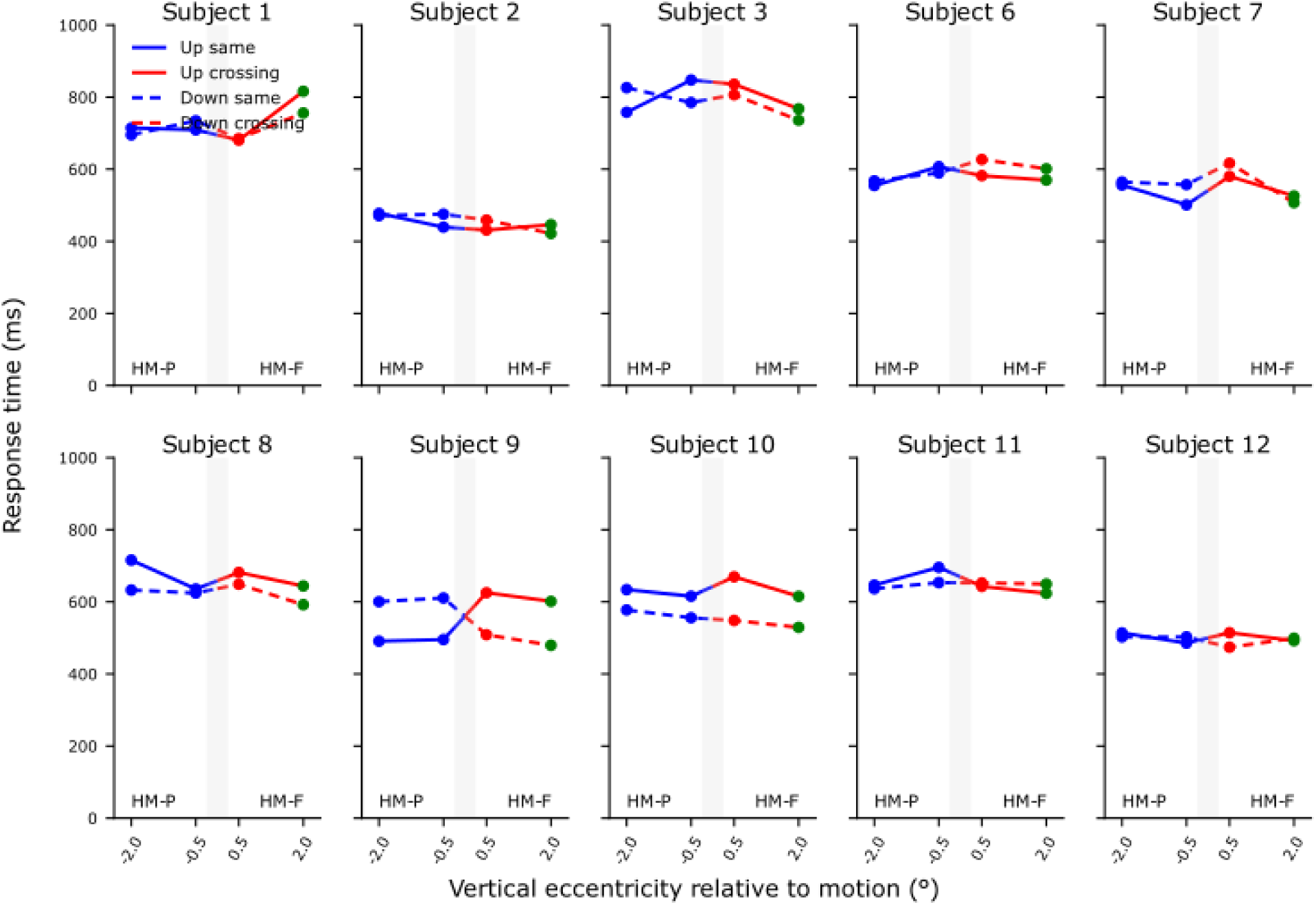
Experiment 3: Flash-lag response time individual results (in ms).

**Supplementary. Figure 9.**
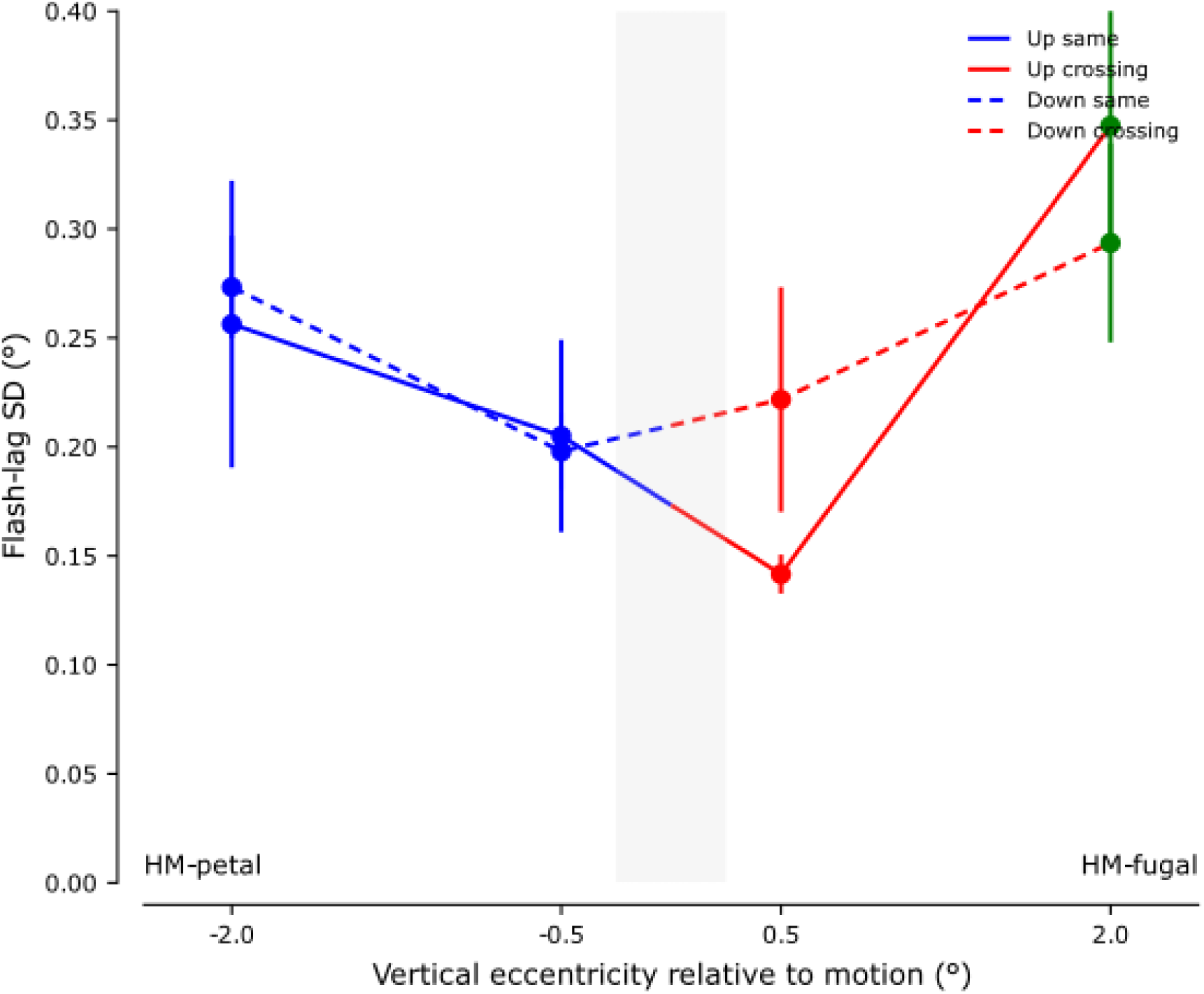
Experiment 3: Average flash-lag standard deviation (in °).

**Supplementary. Figure 10.**
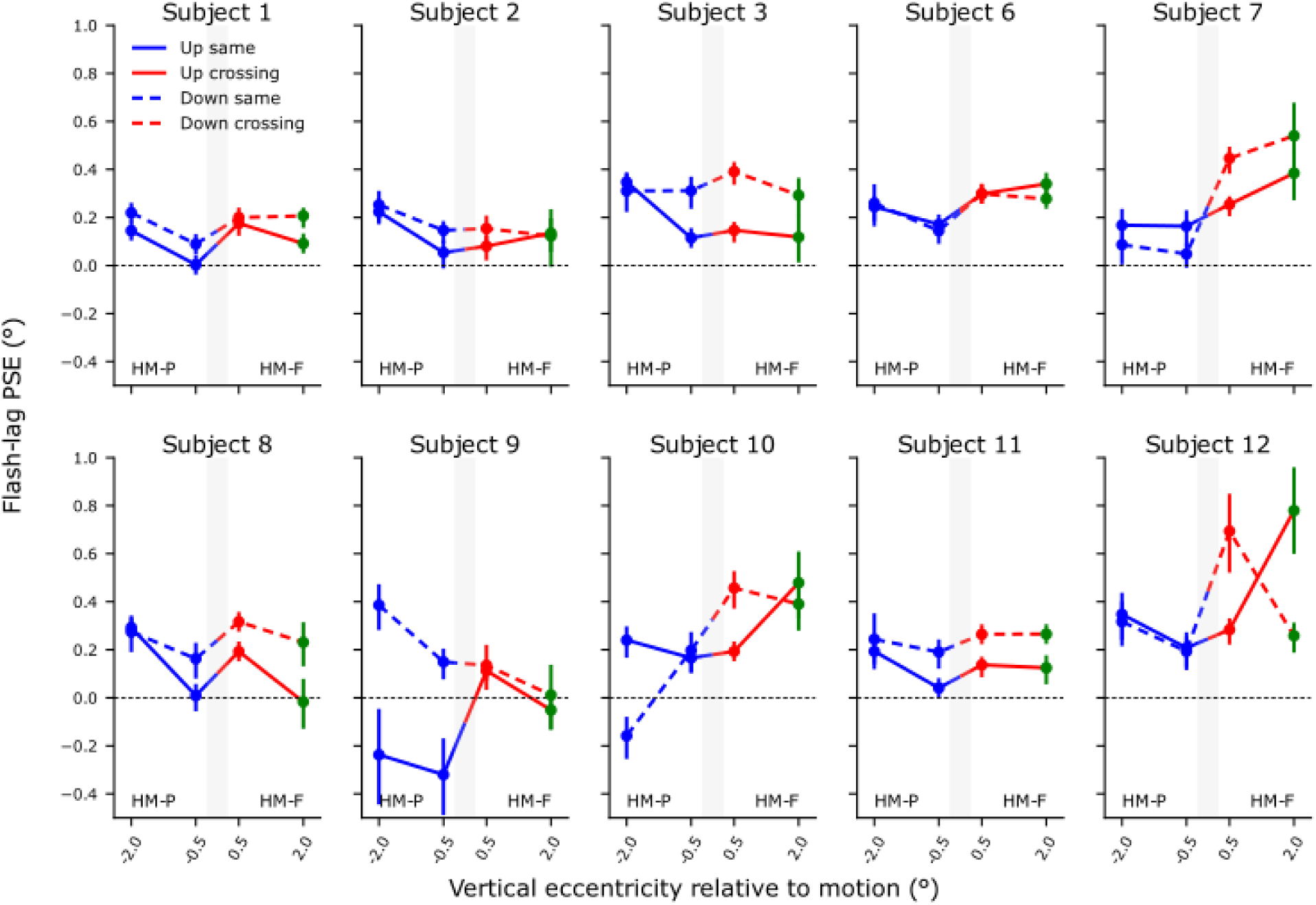
Experiment 3: Flash-lag PSE individual results (in °).

**Supplementary. Figure 11.**
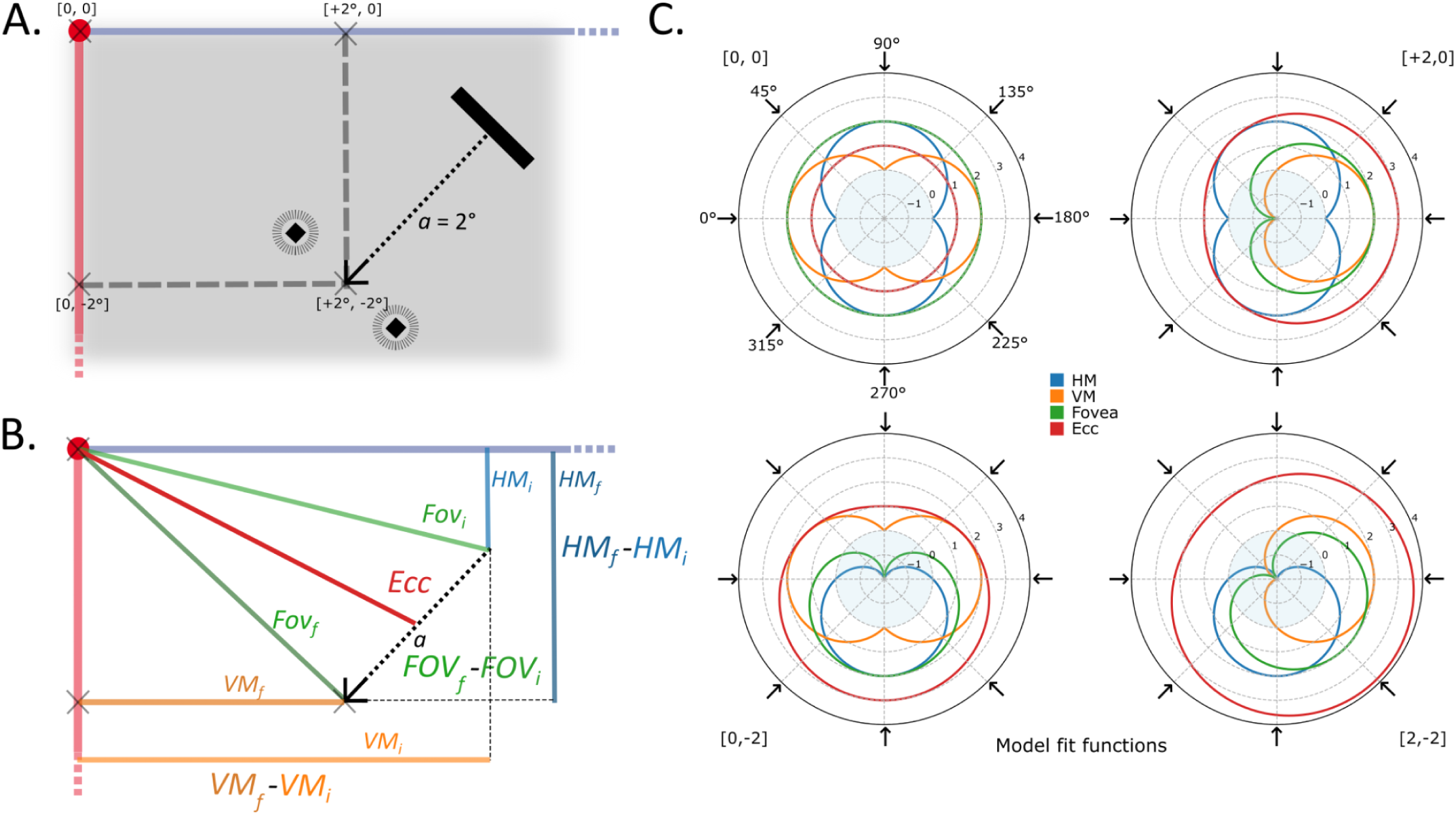
Description of the experiment 4 and illustration of the geometrical modes. **(A)** Schematic of the stimulus used in Experiment 4. Bars could move in 8 possible directions (illustration with a bar moving in the 135° direction). Bars travel a distance of 2° before two flashing targets appear above and below at a certain offset. Bars could travel towards 4 possible positions: [0, 0], [0, -2], [+2, 0], [+2, -2]. **(B)** In the situation illustrated in panel A, the geometrical model is based on the relative distances to each retinotopic features: horizontal meridian (HM in blue), vertical meridian (VM in orange) and fovea (Fov in green), between final (subscript *f*) and starting (subscript *i*) positions. The eccentricity (Ecc in red), measures the foveal distance to the middle of the trajectory. The same color convention as in Figure 1B was used for panel A and B, fovea being represented at [0,0] with a red dot, right HM and lower VM being represented in violet and light red, respectively. **(C)** Polar plots of the geometrical models at each tested position and direction for each of the biases.

**Supplementary. Figure 12.**
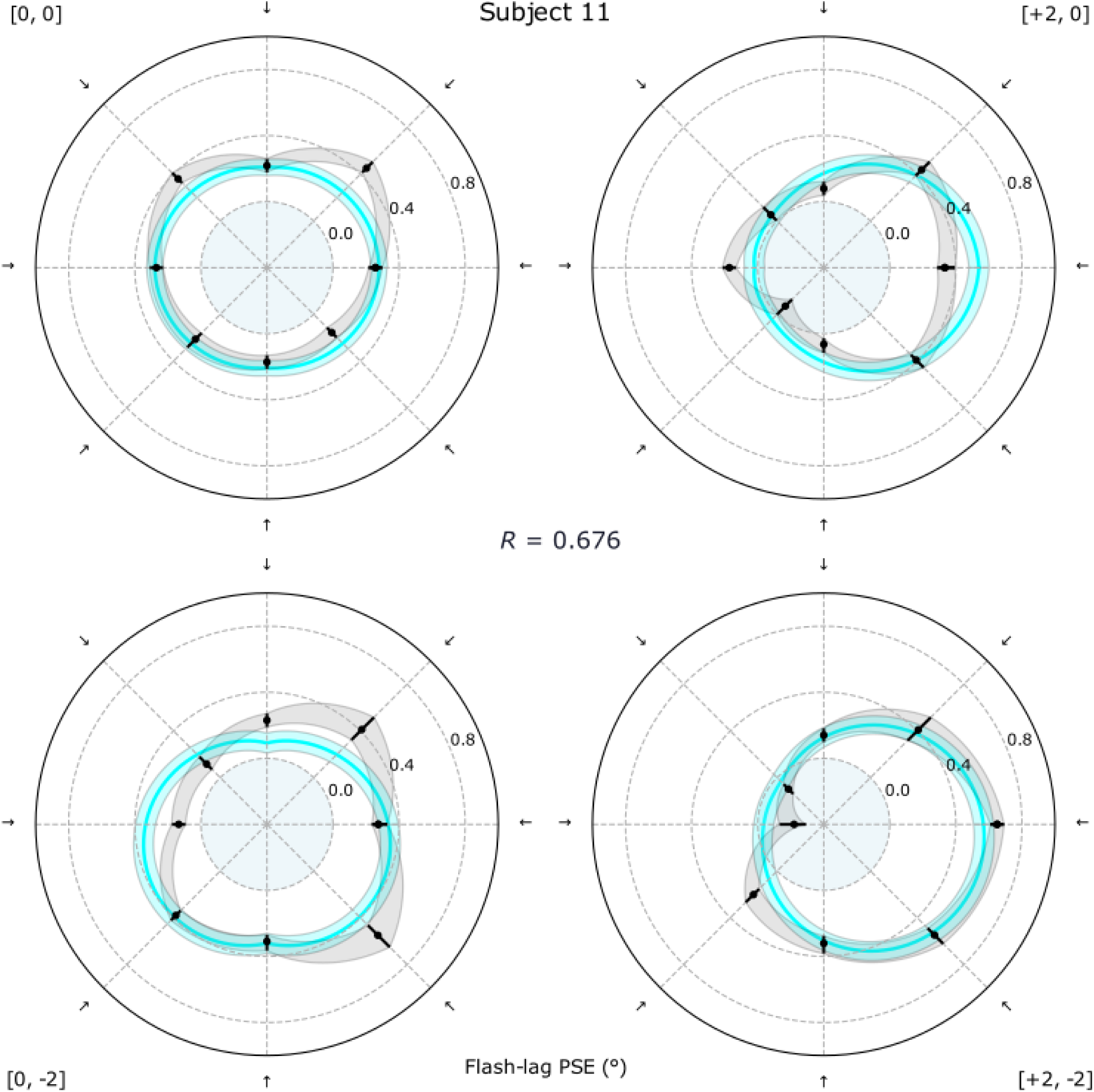
Experiment 4: Typical individual data and fitted model (subject 11).

**Supplementary Figure 13.**
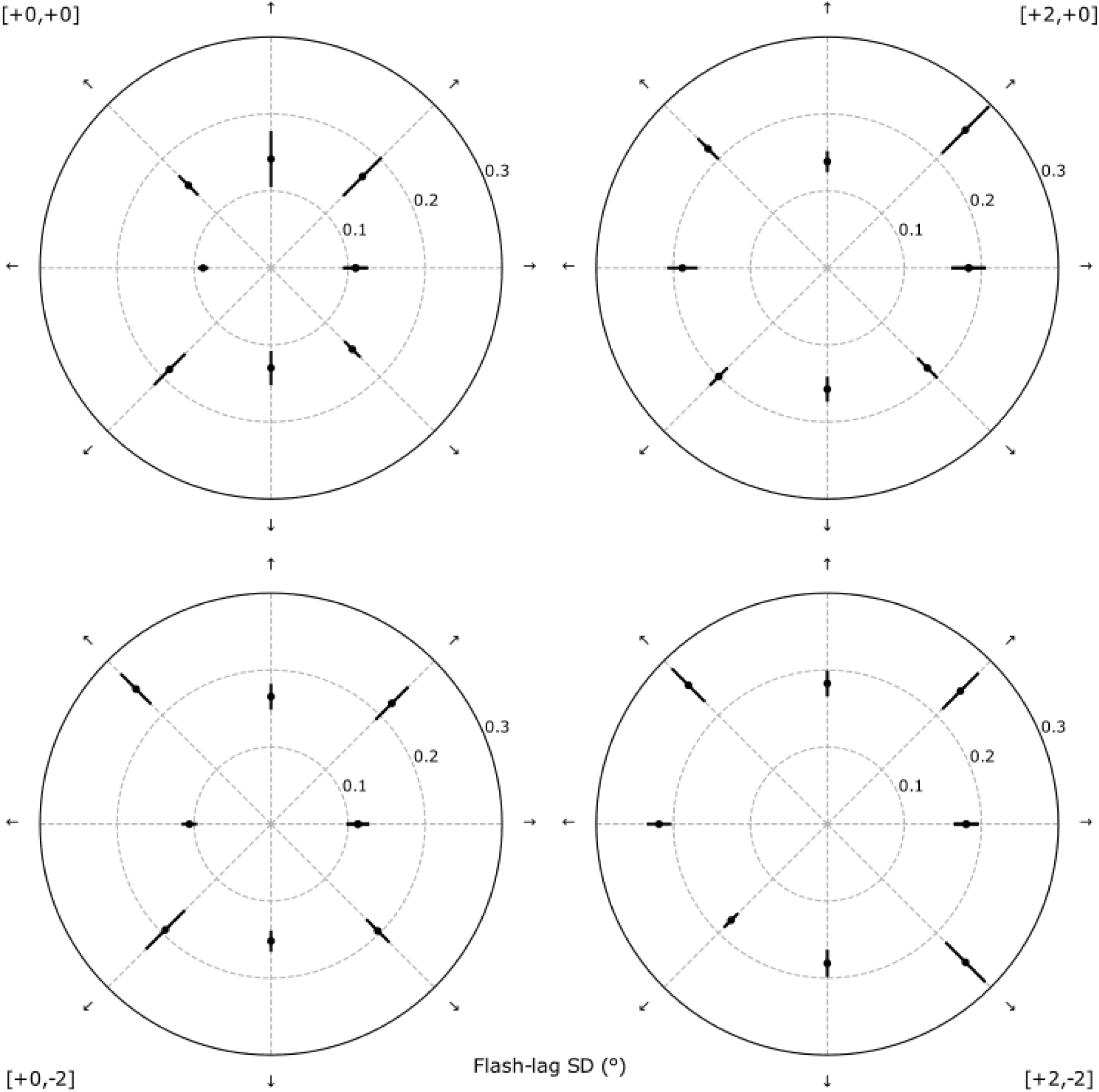
Experiment 4: Average flash-lag standard deviation (in °).

## Bibliography

Amadeo, Maria Bianca, Alessia Tonelli, Claudio Campus, and Monica Gori. 2022. ‘Reduced Flash Lag Illusion in Early Deaf Individuals’. Brain Research 1776 (February): 147744. 10.1016/j.brainres.2021.147744.

Angelucci, Alessandra, and Paul C. Bressloff. 2006. ‘Contribution of Feedforward, Lateral and Feedback Connections to the Classical Receptive Field Center and Extra-Classical Receptive Field Surround of Primate V1 Neurons’. In Progress in Brain Research, edited by S. Martinez-Conde, S. L. Macknik, L. M. Martinez, J. -M. Alonso, and P. U. Tse, vol. 154. Visual Perception. Elsevier. 10.1016/S0079-6123(06)54005-1.

Angelucci, Alessandra, Jonathan B. Levitt, Emma J. S. Walton, Jean-Michel Hupé, Jean Bullier, and Jennifer S. Lund. 2002. ‘Circuits for Local and Global Signal Integration in Primary Visual Cortex’. ARTICLE. Journal of Neuroscience 22 (19): 8633–46. 10.1523/JNEUROSCI.22-19-08633.2002.

Baldo, Marcus V C, Alexandre H Kihara, Janaina Namba, and Stanley A Klein. 2002. ‘Evidence for an Attentional Component of the Perceptual Misalignment between Moving and Flashing Stimuli’. Perception 31 (1): 17–30. 10.1068/p3302.

Benvenuti, Giacomo, Sandrine Chemla, Arjan Boonman, Laurent Perrinet, Guillaume S. Masson, and Frédéric Chavane. 2020. ‘Anticipatory Responses along Motion Trajectories in Awake Monkey Area V1’. Preprint, bioRxiv, April 2. 10.1101/2020.03.26.010017.

Berry, Michael J., Iman H. Brivanlou, Thomas A. Jordan, and Markus Meister. 1999. ‘Anticipation of Moving Stimuli by the Retina’. Nature 398 (6725): 334–38. 10.1038/18678.

Bringuier, Vincent, Frédéric Chavane, Larry Glaeser, and Yves Frégnac. 1999. ‘Horizontal Propagation of Visual Activity in the Synaptic Integration Field of Area 17 Neurons’. Science 283 (5402): 695–99. 10.1126/science.283.5402.695.

Caminiti, Roberto, Filippo Carducci, Claudia Piervincenzi, et al. 2013. ‘Diameter, Length, Speed, and Conduction Delay of Callosal Axons in Macaque Monkeys and Humans: Comparing Data from Histology and Magnetic Resonance Imaging Diffusion Tractography’. Articles. Journal of Neuroscience 33 (36): 14501–11. 10.1523/JNEUROSCI.0761-13.2013.

Caminiti, Roberto, Hassan Ghaziri, Ralf Galuske, Patrick R. Hof, and Giorgio M. Innocenti. 2009. ‘Evolution Amplified Processing with Temporally Dispersed Slow Neuronal Connectivity in Primates’. Proceedings of the National Academy of Sciences 106 (46): 19551–56. 10.1073/pnas.0907655106.

Clarke, Stephanie, and Judit Miklossy. 1990. ‘Occipital Cortex in Man: Organization of Callosal Connections, Related Myelo- and Cytoarchitecture, and Putative Boundaries of Functional Visual Areas’. Journal of Comparative Neurology 298 (2): 188–214. 10.1002/cne.902980205.

Eagleman, David M., and Terrence J. Sejnowski. 2000. ‘Motion Integration and Postdiction in Visual Awareness’. Science 287 (5460): 2036–38. 10.1126/science.287.5460.2036.

Eagleman, David M., and Terrence J. Sejnowski. 2007. ‘Motion Signals Bias Localization Judgments: A Unified Explanation for the Flash-Lag, Flash-Drag, Flash-Jump, and Frohlich Illusions’. Journal of Vision 7 (4): 3. 10.1167/7.4.3.

Essen, DC Van, W. T. Newsome, and J. L. Bixby. 1982. ‘The Pattern of Interhemispheric Connections and Its Relationship to Extrastriate Visual Areas in the Macaque Monkey’. Articles. Journal of Neuroscience 2 (3): 265–83. 10.1523/JNEUROSCI.02-03-00265.1982.

Fröhlich, F. W. 1923. ‘Über Die Messung Der Empfindungszeit’. Zeitschrift Für Sinnesphysiologie 54: 58.

Girard, P., J. M. Hupé, and J. Bullier. 2001. ‘Feedforward and Feedback Connections Between Areas V1 and V2 of the Monkey Have Similar Rapid Conduction Velocities’. Journal of Neurophysiology 85 (3): 1328–31. 10.1152/jn.2001.85.3.1328.

Grinvald, A., E. E. Lieke, R. D. Frostig, and R. Hildesheim. 1994. ‘Cortical Point-Spread Function and Long-Range Lateral Interactions Revealed by Real-Time Optical Imaging of Macaque Monkey Primary Visual Cortex’. Articles. Journal of Neuroscience 14 (5): 2545–68. 10.1523/JNEUROSCI.14-05-02545.1994.

Guo, Kun, Robert G. Robertson, Maribel Pulgarin, et al. 2007. ‘Spatio-Temporal Prediction and Inference by V1 Neurons’. European Journal of Neuroscience 26 (4): 1045–54. 10.1111/j.1460-9568.2007.05712.x.

Harvey, Ben M., and Serge O. Dumoulin. 2011. ‘The Relationship between Cortical Magnification Factor and Population Receptive Field Size in Human Visual Cortex: Constancies in Cortical Architecture’. Articles. Journal of Neuroscience 31 (38): 13604–12. 10.1523/JNEUROSCI.2572-11.2011.

Himmelberg, Marc M., Ekin Tünçok, Jesse Gomez, Kalanit Grill-Spector, Marisa Carrasco, and Jonathan Winawer. 2023. ‘Comparing Retinotopic Maps of Children and Adults Reveals a Late-Stage Change in How V1 Samples the Visual Field’. Nature Communications 14 (1): 1561. 10.1038/s41467-023-37280-8.

Himmelberg, Marc M., Jonathan Winawer, and Marisa Carrasco. 2023. ‘Polar Angle Asymmetries in Visual Perception and Neural Architecture’. Trends in Neurosciences 46 (6): 445–58. 10.1016/j.tins.2023.03.006.

Hogendoorn, Hinze. 2020. ‘Motion Extrapolation in Visual Processing: Lessons from 25 Years of Flash-Lag Debate’. Journal of Neuroscience 40 (30): 5698–705. 10.1523/JNEUROSCI.0275-20.2020.

Hogendoorn, Hinze, and Anthony N. Burkitt. 2018. ‘Predictive Coding of Visual Object Position Ahead of Moving Objects Revealed by Time-Resolved EEG Decoding’. NeuroImage 171 (May): 55–61. 10.1016/j.neuroimage.2017.12.063.

Hubbard, Timothy L. 2014. ‘The Flash-Lag Effect and Related Mislocalizations: Findings, Properties, and Theories’. Psychological Bulletin (US) 140 (1): 308–38. 10.1037/a0032899.

Hubbard, Timothy L. 2020. ‘Representational Gravity: Empirical Findings and Theoretical Implications’. Psychonomic Bulletin & Review 27 (1): 36–55. 10.3758/s13423-019-01660-3.

Hubel, D H, and T N Wiesel. 1967. ‘Cortical and Callosal Connections Concerned with the Vertical Meridian of Visual Fields in the Cat.’ Journal of Neurophysiology 30 (6): 1561–73. 10.1152/jn.1967.30.6.1561.

Ichikawa, Makoto, and Yuko Masakura. 2006. ‘Manual Control of the Visual Stimulus Reduces the Flash-Lag Effect’. Vision Research 46 (14): 2192–203. 10.1016/j.visres.2005.12.021.

Ichikawa, Makoto, and Yuko Masakura. 2010. ‘Reduction of the Flash-Lag Effect in Terms of Active Observation’. *Attention, Perception*, & Psychophysics 72 (4): 1032–44. 10.3758/APP.72.4.1032.

Innocenti, Giorgio M., Kerstin Schmidt, Chantal Milleret, et al. 2022. ‘The Functional Characterization of Callosal Connections’. Progress in Neurobiology 208 (January): 102186. 10.1016/j.pneurobio.2021.102186.

Jancke, Dirk, Wolfram Erlhagen, Gregor Schöner, and Hubert R. Dinse. 2004. ‘Shorter Latencies for Motion Trajectories than for Flashes in Population Responses of Cat Primary Visual Cortex’. The Journal of Physiology 556 (3): 971–82. 10.1113/jphysiol.2003.058941.

Johnson, Philippa Anne, Tessel Blom, Simon van Gaal, Daniel Feuerriegel, Stefan Bode, and Hinze Hogendoorn. 2023. ‘Position Representations of Moving Objects Align with Real-Time Position in the Early Visual Response’. eLife 12 (January): e82424. 10.7554/eLife.82424.

Kanai, Ryota, Bhavin R Sheth, and Shinsuke Shimojo. 2004. ‘Stopping the Motion and Sleuthing the Flash-Lag Effect: Spatial Uncertainty Is the Key to Perceptual Mislocalization’. Vision Research 44 (22): 2605–19. 10.1016/j.visres.2003.10.028.

Kennedy, H., and J. Bullier. 1985. ‘A Double-Labeling Investigation of the Afferent Connectivity to Cortical Areas V1 and V2 of the Macaque Monkey’. Articles. Journal of Neuroscience 5 (10): 2815–30. 10.1523/JNEUROSCI.05-10-02815.1985.

Kennedy, Henry, and Colette Dehay. 1988. ‘Functional Implications of the Anatomical Organization of the Callosal Projections of Visual Areas V1 and V2 in the Macaque Monkey’. Behavioural Brain Research 29 (3): 225–36. 10.1016/0166-4328(88)90027-7.

Kennedy, Henry, Colette Dehay, and Jean Bullier. 1986. ‘Organization of the Callosal Connections of Visual Areas v1 and v2 in the Macaque Monkey’. Journal of Comparative Neurology 247 (3): 398–415. 10.1002/cne.902470309.

Kohler, Peter J., Patrick Cavanagh, and Peter U. Tse. 2017. ‘Motion-Induced Position Shifts Activate Early Visual Cortex’. Frontiers in Neuroscience 11 (April). 10.3389/fnins.2017.00168.

Kontsevich, Leonid L., and Christopher W. Tyler. 1999. ‘Bayesian Adaptive Estimation of Psychometric Slope and Threshold’. Vision Research 39 (16): 2729–37. 10.1016/S0042-6989(98)00285-5.

Lee, Tai Sing, David Mumford, Richard Romero, and Victor A. F. Lamme. 1998. ‘The Role of the Primary Visual Cortex in Higher Level Vision’. Vision Research 38 (15): 2429–54. 10.1016/S0042-6989(97)00464-1.

Li, Hsin-Hung, Won Mok Shim, and Patrick Cavanagh. 2014. ‘Backward Position Shift in Apparent Motion’. Journal of Vision 14 (1): 16. 10.1167/14.1.16.

Linares, Daniel, Joan López-Moliner, and Alan Johnston. 2007. ‘Motion Signal and the Perceived Positions of Moving Objects’. Journal of Vision 7 (7): 1. 10.1167/7.7.1.

Liu, Sirui, Peter U. Tse, and Patrick Cavanagh. 2018. ‘Meridian Interference Reveals Neural Locus of Motion-Induced Position Shifts’. Journal of Neurophysiology 119 (6): 2091–99. 10.1152/jn.00876.2017.

Mackay, D. M. 1958. ‘Perceptual Stability of a Stroboscopically Lit Visual Field Containing Self-Luminous Objects’. Nature 181 (4607): 507–8. 10.1038/181507a0.

Mateeff, S., and J. Hohnsbein. 1988. ‘Perceptual Latencies Are Shorter for Motion towards the Fovea than for Motion Away’. Vision Research 28 (6): 711–19. 10.1016/0042-6989(88)90050-8.

Maus, Gerrit W., Jason Fischer, and David Whitney. 2013. ‘Motion-Dependent Representation of Space in Area MT+’. Neuron 78 (3): 554–62. 10.1016/j.neuron.2013.03.010.

Muller, Lyle, Alexandre Reynaud, Frédéric Chavane, and Alain Destexhe. 2014. ‘The Stimulus-Evoked Population Response in Visual Cortex of Awake Monkey Is a Propagating Wave’. Nature Communications 5. 10.1038/ncomms4675.

Nagai, Masayoshi, Koji Kazai, and Akihiro Yagi. 2002. ‘Larger Forward Memory Displacement in the Direction of Gravity’. Visual Cognition, ahead of print, February 1. world. 10.1080/13506280143000304.

Nijhawan, Romi. 1994. ‘Motion Extrapolation in Catching [9]’. Nature 370 (6487): 256–57. 10.1038/370256B0.

Nijhawan, Romi. 2002. ‘Neural Delays, Visual Motion and the Flash-Lag Effect’. Trends in Cognitive Sciences 6 (9): 387–93. 10.1016/S1364-6613(02)01963-0.

Peirce, Jonathan W. 2007. ‘PsychoPy—Psychophysics Software in Python’. Journal of Neuroscience Methods 162 (1): 8–13. 10.1016/j.jneumeth.2006.11.017.

Reynaud, Alexandre, Guillaume S. Masson, and Frédéric Chavane. 2012. ‘Dynamics of Local Input Normalization Result from Balanced Short-and Long-Range Intracortical Interactions in Area V1’. Journal of Neuroscience 32 (36): 12558–69. 10.1523/JNEUROSCI.1618-12.2012.

Ribeiro, Fernanda Lenita, Ashley York, Elizabeth Zavitz, Steffen Bollmann, Marcello GP Rosa, and Alexander Puckett. 2023. ‘Variability of Visual Field Maps in Human Early Extrastriate Cortex Challenges the Canonical Model of Organization of V2 and V3’. eLife 12 (August): e86439. 10.7554/eLife.86439.

Salin, P. A., J. Bullier, and H. Kennedy. 1989. ‘Convergence and Divergence in the Afferent Projections to Cat Area 17’. Journal of Comparative Neurology 283 (4): 486–512. 10.1002/cne.902830405.

Schira, Mark M., Alex R. Wade, and Christopher W. Tyler. 2007. ‘Two-Dimensional Mapping of the Central and Parafoveal Visual Field to Human Visual Cortex’. Journal of Neurophysiology 97 (6): 4284–95. 10.1152/jn.00972.2006.

Schneider, Marian, Ingo Marquardt, Shubharthi Sengupta, Federico De Martino, and Rainer Goebel. 2019. ‘Motion Displaces Population Receptive Fields in the Direction Opposite to Motion’. Preprint, September 8. 10.1101/759183.

Schwartz, E. L. 1977. ‘Spatial Mapping in the Primate Sensory Projection: Analytic Structure and Relevance to Perception’. Biological Cybernetics 25 (4): 181–94. 10.1007/BF01885636.

Sedigh-Sarvestani, Madineh, and David Fitzpatrick. 2022. ‘What and Where: Location-Dependent Feature Sensitivity as a Canonical Organizing Principle of the Visual System’. Frontiers in Neural Circuits 16. https://www.frontiersin.org/articles/10.3389/fncir.2022.834876.

Shi, Zhuanghua, and Romi Nijhawan. 2008. ‘Behavioral Significance of Motion Direction Causes Anisotropic Flash-Lag, Flash-Drag, Flash-Repulsion, and Movement-Mislocalization Effects’. Journal of Vision 8 (7): 24. 10.1167/8.7.24.

Shi, Zhuanghua, and Romi Nijhawan. 2012. ‘Motion Extrapolation in the Central Fovea’. PLOS ONE 7 (3): e33651. 10.1371/journal.pone.0033651.

Shipp, S., J. D. G. Watson, R. S. J. Frackowiak, and S. Zeri. 1995. ‘Retinotopic Maps in Human Prestriate Visual Cortex: The Demarcation of Areas V2 and V3’. NeuroImage 2 (2, Part A): 125–32. 10.1006/nimg.1995.1015.

Subramaniyan, Manivannan, Alexander S. Ecker, Saumil S. Patel, et al. 2018. ‘Faster Processing of Moving Compared with Flashed Bars in Awake Macaque V1 Provides a Neural Correlate of the Flash Lag Illusion’. Journal of Neurophysiology 120 (5): 2430–52. 10.1152/jn.00792.2017.

Suzuki, Yuta, Sumeyya Atmaca, and Bruno Laeng. 2023. ‘The Lateralized Flash-Lag Illusion: A Psychophysical and Pupillometry Study’. Brain and Cognition 166 (March): 105956. 10.1016/j.bandc.2023.105956.

Tomasi, Simone, Roberto Caminiti, and Giorgio M. Innocenti. 2012. ‘Areal Differences in Diameter and Length of Corticofugal Projections’. Cerebral Cortex 22 (6): 1463–72. 10.1093/cercor/bhs011.

Turner, William, Charlie Sexton, Philippa A. Johnson, Ella Wilson, and Hinze Hogendoorn. 2024. ‘Progressive Multi-Stage Extrapolation of Predictable Motion in Human Visual Cortex’. Preprint, bioRxiv, April 25. 10.1101/2024.04.22.590502.

Turner, William, Charlie Sexton, Philippa A. Johnson, Ella Wilson, and Hinze Hogendoorn. 2025. ‘Predictable Motion Is Progressively Extrapolated across Temporally Distinct Processing Stages in the Human Visual Cortex’. PLOS Biology 23 (5): e3003189. 10.1371/journal.pbio.3003189.

Turner, William, Charlie M. Sexton, and Hinze Hogendoorn. 2023. ‘Neural Mechanisms of Visual Motion Extrapolation’. Preprint, PsyArXiv, September 11. 10.31234/osf.io/tkg3f.

Van Essen, David C., William T. Newsome, and John H. R. Maunsell. 1984. ‘The Visual Field Representation in Striate Cortex of the Macaque Monkey: Asymmetries, Anisotropies, and Individual Variability’. Vision Research 24 (5): 429–48. 10.1016/0042-6989(84)90041-5.

Vater, Christian, Benjamin Wolfe, and Ruth Rosenholtz. 2022. ‘Peripheral Vision in Real-World Tasks: A Systematic Review’. Psychonomic Bulletin & Review 29 (5): 1531–57. 10.3758/s13423-022-02117-w.

Wang, Xi, Alexandre Reynaud, and Robert F. Hess. 2021. ‘The Flash-Lag Effect in Amblyopia’. Investigative Ophthalmology & Visual Science 62 (2): 23. 10.1167/iovs.62.2.23.

Whitney, David, Herbert C. Goltz, Christopher G. Thomas, Joseph S. Gati, Ravi S. Menon, and Melvyn A. Goodale. 2003. ‘Flexible Retinotopy: Motion-Dependent Position Coding in the Visual Cortex’. Science 302 (5646): 878–81. 10.1126/science.1087839.

Yu, H. -H., T. A. Chaplin, and M. G. P. Rosa. 2015. ‘Representation of Central and Peripheral Vision in the Primate Cerebral Cortex: Insights from Studies of the Marmoset Brain’. *Neuroscience Research*, Marmoset Neuroscience, vol. 93 (April): 47–61. 10.1016/j.neures.2014.09.004.

Yu, Hsin-Hao, Declan P. Rowley, Nicholas S. C. Price, Marcello G. P. Rosa, and Elizabeth Zavitz. 2019. ‘A Twisted Visual Field Map in the Primate Cortex Predicted by Topographic Continuity’. Preprint, bioRxiv, November 11. 10.1101/682187.

Zilles, Karl, and Stephanie Clarke. 1997. ‘Architecture, Connectivity, and Transmitter Receptors of Human Extrastriate Visual Cortex’. In Extrastriate Cortex in Primates, edited by Kathleen S. Rockland, Jon H. Kaas, and Alan Peters. Springer US. 10.1007/978-1-4757-9625-4_15.

